# Rapid Identification of Infections Directly from Isolates and Clinical Specimens with the MasSpec Pen Technology

**DOI:** 10.64898/2026.06.19.732454

**Authors:** Manoj Kushwaha, Faith E. Jackobs, Jacob I. Mardick, Coreen L. Johnson, Sarah Bench, Ashley E. Montgomery, James J. Dunn, Rachel D. Downey, Sarmistha B. Hauger, Janne Heirman, Wout Bittremieux, Lindsey M. Kirkpatrick, Livia S. Eberlin

## Abstract

The time-consuming nature of conventional bacterial identification methods, coupled with rising resistance to antibiotics, presents a serious challenge in the rapid diagnosis and treatment of patients suffering from severe infection and sepsis. In this study, we demonstrate that the MasSpec Pen (MSPen) technology enables rapid identification of the most frequently occurring bacterial species in human infections. A total of 755 clinical bacterial isolates prospectively collected from pediatric and adult patients were analyzed using the MSPen to create a library of molecular profiles. The mass spectra obtained presented distinct molecular patterns for bacterial species, with high relative abundances of bacterial metabolites and lipids, including unique quinolone signaling molecules in *Pseudomonas aeruginosa*, phosphoethanolamine (PE) lipids in *Enterobacteriaceae*, and phosphatidylglycerol (PG) lipids in *Staphylococcus* species. Logistic and multinomial lasso models using relative abundances or logarithmic-ratio abundance calculations achieved >95% accuracy for Gram stain typing and >92% for species-level identification in isolates. Random forest models yielded comparable performance, reinforcing the robustness of these findings across approaches. Many of the metabolite and lipid predictive features from each isolate classifier were detected when directly analyzing highly infected synovial fluid, tissue, pleural fluid, and breast aspirate specimens from patients. Notably, significant variations in the natural isotopic abundance of specific lipids were observed in infected patient samples, indicating presence of bacterial infection. This study demonstrates the potential of MSPen for the direct identification of bacterial-specific metabolites and lipids in clinical isolates and specimens, offering promise for future advancements in the MSPen technology for rapid bacterial identification.

**Teaser.:** MSPen shows potential for rapid, direct identification of bacterial metabolites and lipids, enabling accurate bacterial classification.

## Introduction

The global escalation of antimicrobial-resistant bacterial infections, coupled with inadequate diagnostic pipelines for timely diagnosis and ineffective treatment options, represents one of the most urgent threats to humanity (*1*). Antimicrobial-resistant bacterial infections are expected to claim up to 8.2 million lives annually by 2050, nearly doubling current estimates of 4.71 million annual deaths. In 2019 alone, six bacterial species, *Staphylococcus aureus, Escherichia coli, Streptococcus pneumoniae, Klebsiella pneumoniae, Pseudomonas aeruginosa*, and *Acinetobacter baumannii* accounted for 3.57 million of the total 7.7 million deaths associated with bacterial infections worldwide (*2, 3*). Many of these species are members of the normal commensal bacterial flora of specific organs and tissues; however, under conditions such as immune compromise, dysbiosis, or colonization of non-native anatomical sites, they can become opportunistic pathogens (*4*). Alarmingly, pathogenic bacteria employ diverse mechanisms to acquire and disseminate antimicrobial resistance, resulting in varying resistance profiles across species and strains. For example, Methicillin-resistant *S. aureus* (MRSA) is of particular concern as these strains are resistant to β-lactam antibiotics, the standard treatment for routine *Staphylococcus* infections, and were responsible for over 100,000 deaths in 2019 (*2, 3, 5, 6*).

The current gold standard for bacterial identification relies on phenotypic methods, including culture on selective media, colony enrichment, biochemical testing, and phenotypic antimicrobial susceptibility testing (AST) (*7-9*). These approaches typically require two to five days to yield definitive results, and even longer for fungi and fastidious pathogens. Delayed pathogen identification may hinder timely optimization of therapy, contributing to prolonged hospitalization, increased risk of septic shock or mortality, and unnecessary continuation of inappropriate antimicrobial treatment, which carries significant downstream risks including *Clostridioides difficile* infection, secondary infections, and antimicrobial resistance. Given these limitations, improving the breadth, speed, and accuracy of bacterial diagnostics is a critical priority. Nucleic acid amplification technologies have been increasingly incorporated into microbiology workflows for bacterial detection and identification in cases where specimens may be difficult to culture, such as cerebrospinal fluid and synovial fluid. However, these methods have limitations in terms of sensitivity and specificity and are often utilized when microbial infection cannot be ascertained by other methods (*10*). Mass spectrometry (MS) techniques have been explored for decades as tools for bacterial identification (*11, 12*). Matrix-assisted laser desorption/ionization time-of-flight MS (MALDI-TOF MS) has become an essential clinical technology for rapid identification of bacterial species through detection of protein markers that are characteristic of pathogenic microorganisms (*13*). MALDI-TOF MS can also aid in antimicrobial resistance profiling through the detection of penicillin-binding proteins and β-lactamase activity(*14*). In the research setting, many alternative MS-based approaches are also being developed for bacteria identification (*15–18*). Liquid chromatography tandem mass spectrometry (LC–MS/MS), for example, has been applied to metabolomic flux analysis of common blood-borne pathogens, enabling identification of bacterial infections within ∼10 hours (*19, 20*). Alternatively, Chen *et al.* recently employed rapid evaporative ionization MS (REIMS) to directly analyze clinical isolates and colorectal tissues (*21*). In the study, the authors identified 359 taxon-specific markers in clinical isolates, primarily metabolites and lipids, many of which were also detected in infected colorectal tissue, highlighting the potential value of direct MS analysis of small molecules for bacteria identification.

We have developed the MasSpec Pen (MSPen) technology for direct, non-destructive analysis of tissues and fluids (*22*). The MSPen consists of a handheld device connected to an interface and a mass spectrometer. When the device is placed in contact with a tissue, a solvent droplet is delivered to the tip, enabling gentle extraction of molecules from the tissue surface. The droplet is then aspirated into a heated interface integrated to a mass spectrometer, where analytes are rapidly vaporized for MS analysis (*23*). The resulting mass spectra capture a rich array of metabolites and lipids that can be leveraged to build robust statistical classifiers (*24, 25*). We have successfully applied this workflow to cancer detection and intraoperative surgical margin assessment (*22, 26–28*). We have also shown in a pilot study that MSPen-based analysis can discriminate bacterial species in banked standard culture samples and identify bacterial molecules in infected clinical specimens (*29*). Building on this work, here we applied the MSPen to analyze 755 bacterial clinical isolates and 36 pediatric and adult tissue and fluid specimens prospectively collected from clinical practice. We identified metabolites and lipids that serve as bacteria-specific markers for Gram typing, genus-level discrimination between *Staphylococcus* and *Streptococcus*, differentiation of Group A (GAS) and Group B *Streptococcus* (GAS), and species-level identification. The statistical models built from the molecular data achieved over 92% accuracy for species-level classification on independent pure isolate testing sets, and enabled identification of predictive molecular features associated with each species. Detection of these molecules directly from clinical specimens highlights the potential of MSPen for direct, rapid detection of the most frequently encountered bacterial pathogens.

## Results

### Molecular analysis and molecular trends of clinical isolates

A total of 755 pure bacterial isolates were analyzed using the MSPen, including *Enterobacter cloacae* complex (n = 40), *E. coli* (n = 45), *Haemophilus influenzae* (n = 41), *K. pneumoniae* (n = 50), *P. aeruginosa* (n = 81), *Enterococcus faecalis* (n = 13), *Enterococcus faecium* (n = 14), *Staphylococcus epidermidis* (n = 48), *S. aureus* (including MRSA, n = 158, and methicillin-sensitive [MSSA], n = 170), *Streptococcus pyogenes* (GAS, n = 40), *Streptococcus agalactiae* (GBS, n = 27), and *S. pneumoniae* (n = 28) (**Fig. 1A**). We qualitatively observed that the negative ion mode mass spectra from each bacterial species exhibited characteristic molecular profiles, primarily consisting of metabolites and lipids. Across all isolates, amino acids, dipeptides, short-chain fatty acids (FA), and quorum-sensing metabolites were detected within the *m/z* 100 - 500 range, while more complex lipid species, including ceramides (Cer), phosphatidylglycerols (PG), and phosphatidylethanolamines (PE) were detected within the *m/z* 650 - 800 range (**Fig. 1B**). Molecular identities were tentatively assigned to 214 ions detected based on high mass accuracy measurements, of which 110 were further identified using MS/MS (**Supplementary Data Excel Sheet 1**). For visualization of molecular similarities and global trends among isolates, t-distributed stochastic neighbor embedding (t-SNE) was applied to all principal components (PCs) derived from principal components analysis (PCA). Consistent with the mass spectral trends observed, the resulting t-SNE projection revealed distinct clustering patterns that broadly reflected species-level classification of the bacterial isolates (**Fig. 1C**). Capitalizing on these molecular differences, statistical classification of clinical isolates was performed using a variety of statistical approaches (Methods), and the results obtained for each model developed are described next.

**Figure 1.**
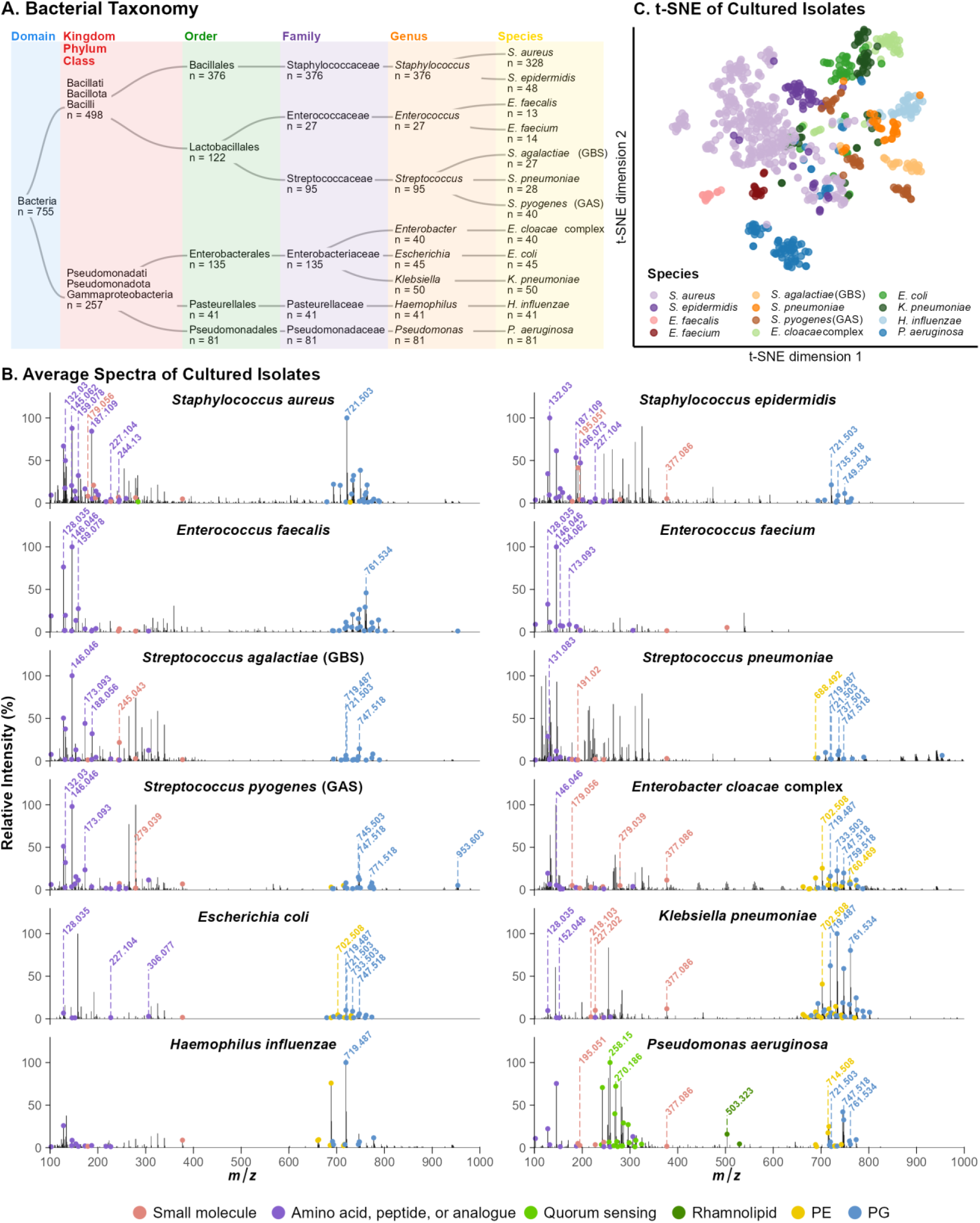
**(A)** Taxonomy of bacterial isolates and the number of each (n) analyzed by MSPen. **(B)** t-SNE plot visualization of MSPen spectra colored by genera; points that are close together have similar mass spectra. **(C)** Average MSPen spectra of bacterial species with molecules of interest annotated by colored circles.

### Gram Stain Classification

Gram staining is a critical first step in bacterial identification, providing essential diagnostic information to guide early treatment decisions. Of the 755 clinical isolates, 257 were Gram-negative and 498 were Gram-positive bacterium. Initial molecular assessment of Gram stain classification within the *m/z* 650 – *m/z* 800 lipid range revealed prominent PE and PG peaks with distinct patterns of relative abundance between bacterial groups. In the mass spectra from Gram negative members of the *Enterobacteriaceae* family, including *E. coli*, *K. pneumoniae*, and the *E. cloacae* complex*, m/z* 702.508 (PE 33:1) and *m/z* 719.487 (PG 32:1) exhibited higher relative abundance compared to the mass spectra of Gram-positive species **(Fig. 1B)**. In contrast, Gram-positive *Staphylococcus* species displayed higher relative abundance of *m/z* 721.503 (PG 32:0), particularly in *S. aureus* and *S. epidermidis*. Among *Streptococcus* species (*S. pyogenes, S. agalactiae, and S. pneumoniae*), *m/z* 747.518 (PG 34:1) was detected at consistently higher relative abundance than in Gram-negative bacteria **(Fig. 1B)**. To evaluate Gram stain classification, logistic regression with lasso penalization was applied to predict Gram-positive versus Gram-negative class membership by first using the normalized relative abundances of the ions, hereby referred to as “relative intensity lasso”. The resulting model achieved a cross-validation (CV) overall accuracy of 98.3% on the training set (n = 525) and 100.0% on the test set (n = 230) (**Fig. S1).** A total of 38 molecular features were selected for Gram stain prediction **(Fig. 2A)**. Features predictive of Gram-negative classification included *m/z* 258.150 (2-heptyl-4-hydroxyquinoline N-oxide), *m/z* 279.039 (pseudouridine), *m/z* 714.508 (PE 34:2) and *m/z* 719.487 (PG 32:1). In contrast, features predictive of Gram-positive classification included *m/z* 159.078 (alanyl-alanine), *m/z* 173.093 (glycyl-valine) and *m/z* 688.492 (PE 32:1).

**Figure 2.**
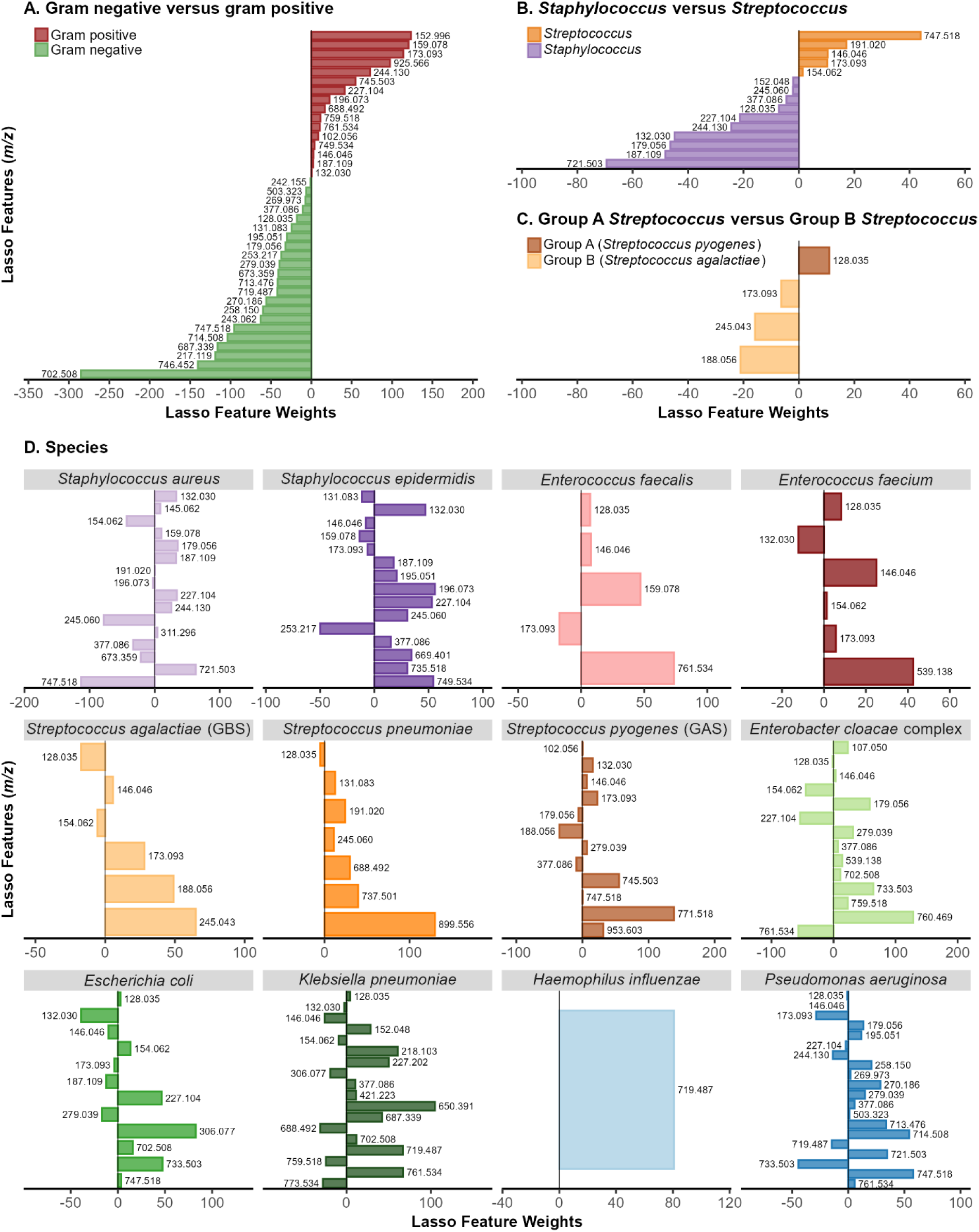
Bar plots showing Lasso feature weights for different levels of bacterial taxonomic classification: (A) Gram stain, (B) *Staphylococcus* vs. *Streptococcus*, (C) Group A vs. Group B *Streptococcus,* and (D) species. In panels (A-C), positive feature weights are associated with one class and negative feature weights are associated with the other class. In species bar plots, positive feature weights indicate contribution of that molecular feature toward predicting that species, whereas negative weights indicate molecular features that detract from prediction of that species. These features correspond to *m*/*z* values tentatively assigned using high-resolution mass spectrometry (±5 ppm mass accuracy) and tandem mass spectrometry (±10 ppm mass accuracy). Their tentative identity, chemical formulas and ionic forms are provided in Supplementary Data, Excel Sheet 1.

Given the variability in lipid peak intensities across pure isolates and observed patterns in the ratios of specific pairs of lipid species, we next applied a lasso framework using pairwise logarithmic ratios of molecular features, hereby referred to as “log-ratio lasso” (**Methods**). This approach achieved a CV accuracy of 97.9% on the training set and 99.6% on the test set for Gram stain classification (**Fig. S2)**. A total of 30 predictive ratio features were selected, with 10 associated with gram-positive classification and 20 with Gram-negative classification. Consistent with relative abundance lasso results, log-ratio lasso identified *m/z* 196.073 (acetyl-histidine) and *m/z* 244.130 (asparaginyl-leucine) as features associated with Gram-positive bacteria, while *m/z* 242.155 (2-heptyl-4-quinolone [HHQ]), *m/z* 749.534 (PG 34:0), *m/z* 728.524 (PE 35:2) and *m/z* 735.518 (PG 33:0) were associated with Gram-negative classification. Ratios of *m/z* 146.046 to *m/z* 714.508 and *m/z* 196.073 to *m/*z 377.086 were predictive of Gram-positive bacteria, whereas the ratios of *m/z* 242.155 to *m/z* 735.518 and *m/z* 714.508 to *m/z* 737.497 were predictive of Gram-negative bacteria. Importantly, strong concordance was observed between the relative abundance lasso and log-ratio lasso models in the selection of molecular features relevant for Gram stain classification. Random forest classification was also performed to determine whether ensemble methods may improve predictive accuracies. The resulting model obtained 99.6% overall accuracy on the test set (**Fig.S3**).

### Staphylococcus and Streptococcus Genera Classification

*Staphylococcus and Streptococcus* are among the most frequently encountered and clinically significant Gram-positive pathogens. Accurate discrimination between these genera is essential because of their distinct antimicrobial resistance profiles, pathogenic mechanisms, and implications for appropriate therapeutic selection. The clinical isolate cohort included *Staphylococcus* species (*S. epidermidis*, n = 48; *S. aureus*, including MRSA, n = 158, and MSSA, n = 170) and *Streptococcus* species (*S. pyogenes*, n = 40; *S. agalactiae*, n = 27; and *S. pneumoniae*, n = 28), comprising a total of 376 *Staphylococcus* species isolates and 95 *Streptococcus* species isolates.

Logistic regression with relative intensity lasso achieved a CV accuracy of 99.7% on the training set and 98.6% on the test set (**Fig. S4**). The model selected 15 molecular features that were predictive of genus-level classification. Features predictive of *Staphylococcus* genus included *m/z* 132.030 (aspartic acid), *m/z* 179.056 (aldohexose), and *m/z* 721.503 (PG 32:0). In contrast, features predictive of the *Streptococcus* genus included *m/z* 146.046 (glutamic acid), *m/z* 173.093 (glycyl-valine), and *m/z* 747.518 (PG 34:1) (**Fig. 2B**). Logistic regression with the log-ratio lasso model achieved a CV accuracy of 98.5% on the training set and 100.0% on the test set (**Fig. S5**). The model selected 18 predictive ratio features for classification. Selected ratios predictive of the *Streptococcus* genus include *m/z* 245.043 (glycerophosphoglycerol) to *m/z* 763.549 (PG 35:0) and *m/z* 745.503 (PG 34:2) to *m/z* 763.549. In contrast, selected ratios predictive of the *Staphylococcus* genus include the ratios of *m/z* 187.109 (N-acetyl-lysine) to *m/z* 245.043 (glycerophosphoglycerol) and *m/z* 191.020 (2-dehydro-3-deoxy-glucaric acid) to *m/z* 717.471 (PG 32:2). Random forest classification similarly achieved 100.0% overall accuracy on the test set (**Fig. S6**).

### GAS versus GBS Classification

GAS and GBS are clinically important gram-positive pathogens. GBS primarily affects neonates and immunocompromised individuals, whereas GAS is associated with a broad spectrum of diseases ranging from pharyngitis to severe invasive diseases. Accurate discrimination between GAS and GBS is clinically important due to differences in disease presentation and antimicrobial susceptibility profiles, including the presence of β-lactamase-mediated resistance in certain strains, which can influence therapeutic decision making.

We performed MSPen analysis of *S. pyogenes* (GAS, n = 40) and *S. agalactiae* (GBS, n = 27) isolates. Logistic regression with relative intensity lasso was used to differentiate GAS and GBS at the species level. The model achieved a CV overall accuracy of 97.9% on the training set and 90.0% accuracy on the test set (**Fig. S7**) using a total of 4 predictive features. The one feature predictive of GAS was *m/z* 128.035 (C_5_H_7_NO_3_), whereas features predictive of GBS included *m/z* 173.093 (glycyl-valine), *m/z* 188.056 (N-acetylglutamate) and *m/z* 245.043 (glycerophosphoglycerol) **(Fig. 2C)**. Logistic regression with the log-ratio lasso model achieved 100.0% accuracy on both the training and test sets with 5 predictive ratio features, all of which were weighted toward the GBS class (**Fig. S8**). The predictive ratio features included *m/z* 188.057 (N-acetylglutamate) to *m/z* 731.487 (PG 33:2) and *m/*z 188.057 to *m/*z 771.518 (PG 36:3) toward GBS. Similarly, random forest classification achieved 100.0% accuracy on the test set.

### Bacterial Species Classification

Accurate identification of individual bacterial species is critical for guiding appropriate antimicrobial therapy, as antibiotic efficacy varies substantially across microbial taxa. Clinical isolates of a total of 12 species representing *E. cloacae* complex (n = 40), *E. coli* (n = 45), *H. influenzae* (n = 41), *K. pneumoniae* (n = 50), *P. aeruginosa* (n = 81), *E. faecalis* (n = 13), *E. faecium* (n = 14), *S. epidermidis* (n = 48), *S. aureus* (MRSA, n = 158; MSSA, n = 170), *S. pyogenes* (GAS, n = 40), *S. agalactiae* (GBS, n = 27), *and S. pneumoniae* (n = 28) were analyzed to evaluate the ability to classify bacteria per species. A multinomial classification model was first built using relative intensity lasso, achieving a CV species-specific classification overall accuracy of 93.9% in the training set and 96.1% overall accuracy in the test set (**Table 1**). In total, 57 predictive features were selected by the model for species-level classification (**Fig. 2D**).

**Table 1.** Logistic regression Lasso-based species classification results on the training and test set.

| Species | Training Set |  | Test Set |  |
| --- | --- | --- | --- | --- |
|  | Recall Rate | Correct/Total | Recall Rate | Correct/Total |
| *E. coli* | 83.87% | 24/31 | 92.86% | 13/14 |
| *E. cloacae* complex | 85.71% | 24/28 | 100.00% | 12/12 |
| *E. faecalis* | 88.89% | 8/9 | 100.00% | 4/4 |
| *E. faecium* | 90.00% | 9/10 | 100.00% | 4/4 |
| *H. influenzae* | 100.00% | 29/29 | 100.00% | 12/12 |
| *K. pneumoniae* | 88.57% | 31/35 | 100.00% | 15/15 |
| *P. aeruginosa* | 89.47% | 51/57 | 100.00% | 24/24 |
| *S. aureus* | 98.67% | 223/226 | 96.94% | 99/102 |
| *S. epidermidis* | 87.88% | 29/33 | 80.00% | 12/15 |
| *S. agalactiae* | 100.00% | 19/19 | 87.50% | 7/8 |
| *S. pneumoniae* | 80.00% | 16/20 | 87.50% | 7/8 |
| *S. pyogenes* | 100.00% | 28/28 | 100.00% | 12/12 |
| Overall Accuracy | 93.90% | 493/525 | 96.09% | 221/230 |

Within the Gram-positive *Staphylococcus* species analyzed, commonly detected features included *m/z* 132.030 (aspartic acid), *m/z* 187.109 (N-acetyl-lysine), *m/z* 227.104 (hydroxyprolyl-proline), and *m/*z 377.086 (lactose) **(Fig. 2D)**. These detected features were all weighted towards the *Staphylococcus* group of species, except for *m/z* 377.086 which was predictive of *S. epidermidis* and was weighted against *S. aureus*. Additional predictive features for *S. aureus* included *m/z* 244.130 (asparaginyl-leucine) and *m/z* 721.503 (PG 32:0). In contrast *m/z* 749.534 (PG 34:0) was weighted towards *S. epidermidis*.

Species-specific lasso weighting for the *Streptococcus* genus (*S. pneumoniae*, *S. pyogenes*, and *S. agalactiae*) indicated predictive features for *S. pneumoniae* including *m/z* 688.492 (PE 32:1), *m/*z 737.501 (PG 32:0;O), and *m/z* 899.556 (PG 43:6), for *S. pyogenes* with *m/*z 745.503 (PG 34:2), m/z 771.518 (PG 36:3), and *m/z* 953.602, and for *S. agalactiae* with *m/z* 173.093 (glycyl-valine), m/z 188.056 (N-acetylglutamate), and *m/*z 245.043 (glycerophosphoglycerol) (**Fig. 2D)**. The lipids at *m/z* 688.492, *m/z* 745.503, and *m/z* 771.518 were commonly detected in the mass spectra of each of the *Streptococcus* species.

Within the *Enterococcus* genus, *E. faecium* and *E. faecalis* shared similar mass spectral profiles, including common peaks at *m/z* 747.522 (PG 34:1) and *m/z* 761.534 (PG 35:1), which were detected at different relative abundances. Commonly predictive features for both species included *m/z* 128.035 (C_5_H_7_NO_3_) and *m/z* 146.046 (glutamic acid). Features weighted towards *E. faecium* included *m/*z 173.093 (glycyl-valine) and *m/z* 539.139 (raffinose), while features weighted towards *E. faecalis* included *m/z* 159.078 (alanyl-alanine) and *m/z* 761.534 (PG 35:1) (**Fig. 2D)**.

Molecular profiles of the *Enterobacteriaceae* family (*E. coli, E. cloacae* complex*, and K. pneumoniae*) shared common lipid features at *m/z* 702.508 (PE 33:1) and *m/z* 733.503 (PG 33:1). Species-specific features weighted towards *E. coli* included *m/z* 154.062 (histidine) and *m/z* 733.503 (PG 33:1), while features weighted towards *E. cloacae* complex included *m/z* 279.039 (pseudouridine) and *m/z* 759.518 (PG 35:2), and towards *K. pneumoniae* included *m/z* 719.487 (PG 32:1) and *m/z* 760.469 (PE 35:4) (**Fig. 2D**). The *H. influenzae* species consisted of one lone lasso predictive feature at *m/z* 719.487 (PG 32:1) (**Fig. 2D)**.

The mass spectra from *P. aeruginosa* presented a unique metabolic profile characterized by quinolone signaling molecules that were specific to this species. These included *m/z* 258.150 (2-heptyl-4-hydroxyquinoline N-oxide [HQNO]) and *m/z* 270.186 (2-nonyl-4-hydroxyquinoline [NHQ]) (**Fig. 2D and Fig. S10**). In total, 29 2-alkyl-quinolone class molecules were identified in *P. aeruginosa* isolates (**Supplementary Data Excel Sheet 2**). In addition to quinolones, lipid species detected in *P. aeruginosa* included *m/*z 719.487 (PG 32:1), *m/z* 721.503 (PG 32:0), and *m/z* 747.518 (PG 34:1) **(Fig. S11)**, among others.

We also applied multinomial log-ratio lasso and random forest to the same dataset to further explore species-level predictability of the dataset with different statistical methods. Log-ratio lasso achieved an overall CV accuracy of 92.95% on the training set and 94.35% on the test set (**Fig. S12 and S13)** using a total of 284 ratio features. The multinomial random forest classifier was also trained on the same dataset, yielding accuracies of 94.6% on the training set and 96.8% on the test set (**Fig. S14**). Collectively, species-level classification using relative abundance multinomial lasso, multinomial log-ratio lasso, and multinomial random forest models achieved accuracies ranging from 92% to 97%.

### Methicillin-Resistant versus Methicillin-Sensitive *S. aureus*

For clinical samples identified with *S. aureus* infection, discrimination between MRSA and MSSA is required to evaluate effective target antibiotics for treatment. In our dataset of 328 *S. aureus* isolates, a total of 170 were MSSA while 158 were MRSA. Qualitative evaluation of the MSPen mass spectra profiles of MRSA and MSSA revealed similar profiles of metabolites and lipid peaks that are characteristic of *S. aureus*, with small variations in the relative abundances of *m/z* 707.488 (PG 31:0), *m/z* 731.487 (PG 33:2), *m/z* 733.503 (PG 33:1), and *m/z* 735.518 (PG 33:0) (**Fig. S15**). Despite the similarities in mass spectra, a similar strategy was applied to attempt to classify the isolates as MSSA and MRSA using logistic regression with relative abundance lasso. The resulting lasso model achieved a classification accuracy of 67.3% on the training set and 63.7% on the test set. Similar performance was observed for the log-ratio lasso model and with random forests, where accuracies of 62.4% and 61.8% were obtained using log-ratio lasso on the training and test sets, respectively (**Fig. S16 and S17**). With random forest, an out-of-fold cross-validation accuracy of 64.7% was achieved and 65.7% test set accuracy (**Fig. S18**). Predictive features for the relative intensity and log-ratio lasso models are provided in **Supplementary Data Excel Sheets 3 and 4**. Within the features detected by relative intensity lasso, *m/z* 572.482 (Cer 34:1), *m/z* 707.487 (PG 31:0), *m/z* 731.487 (PG 31:2) were weighted towards MRSA, while *m/z* 719.487 (PG 32:1), *m/z* 733.503 (PG 33:1) and *m/z* 759.518 (PG 35:2) were weighted towards MSSA.

### Identification of Bacterial Infection in Clinical Specimens

We next evaluated whether bacterial markers identified in clinical cultured isolates could be detected directly in infected patient specimens. In total, 39 infected specimens (n = 36 monomicrobial, n = 3 polymicrobial) with abundant bacterial growth (3+ and 4+ bacterial growth on solid media plating) of at least one bacterial species based on clinical microbiology assessment were analyzed (**Supplementary Data Excel Sheet 5**). The clinical cohort included a diverse set of infected tissues and fluids, including abscesses of the prostate, breast, thigh, axilla, and prepatellar region, prosthetic joint infections of the knee and hip, wound and ulcer tissues, bursa and lymph node aspirates, lung and prostate tissues, and synovial, pleural, thoracic, and cerebrospinal fluids.

A total of 19 monomicrobial patient specimens analyzed with the MSPen were infected with *S. aureus*, including a sample of breast abscess (sample 1046) and a sample of infected right elbow bursa fluid (sample 1275), both of which exhibited 4+ (abundant) *S. aureus* growth. Direct comparison of MSPen mass spectra acquired from infected breast abscess and elbow bursa fluid specimens and their corresponding *S. aureus* isolates revealed overlap in the detection of several molecular features. As shown in **Fig. 3A**, characteristic lipid peaks at *m/z* 693.472 (PG 30:0) and *m/z* 721.503 (PG 32:0) were detected in the infected breast abscess, while the peaks at *m/z* 735.518 (PG 33:0) and *m/z* 749.534 (PG 34:0) were detected in the infected bursa fluid. The metabolite ions at *m/z* 132.030 (aspartic acid), *m/*z 159.078 (alanyl-alanine), *m/*z 179.056 (aldohexose), *m/*z 227.104 (hydroxyprolyl-proline), and *m/z* 311.296 (FA 20:0) were also observed in the infected specimens and their respective isolates (**Fig. 3A**). As we previously described, these ions were consistently detected in the mass spectra of clinical isolate of several *Staphylococcus* species, including *S. epidermidis and S. aureus*, and were selected in our classifiers as predictive for these species (**Figs. 1B, 2D**). The mass spectra of two clinical specimens infected with *Streptococcus* species also presented lipid profiles that were characteristic of *Streptococcus* infection. For example, the MSPen mass spectra of a sample of thigh abscess and a sample of synovial fluid infected with *Streptococcus* revealed characteristic lipid ions at *m/*z 745.503 (PG 34:2), *m/*z 747.518 (PG 34:1), *m/*z 771.518 (PG 36:3), and *m/*z 773.534 (PG 36:2) (**Fig. S19**). Although these ions are also detected in human tissues, they were also consistently detected in the studied *Streptococcus* species and matched those observed in isolate mass spectra. Importantly, changes in the relative abundance of these lipid ions and their isotopic ratios were observed in infected human specimens as an indication of infection, which we further describe in the next section.

**Figure 3.**
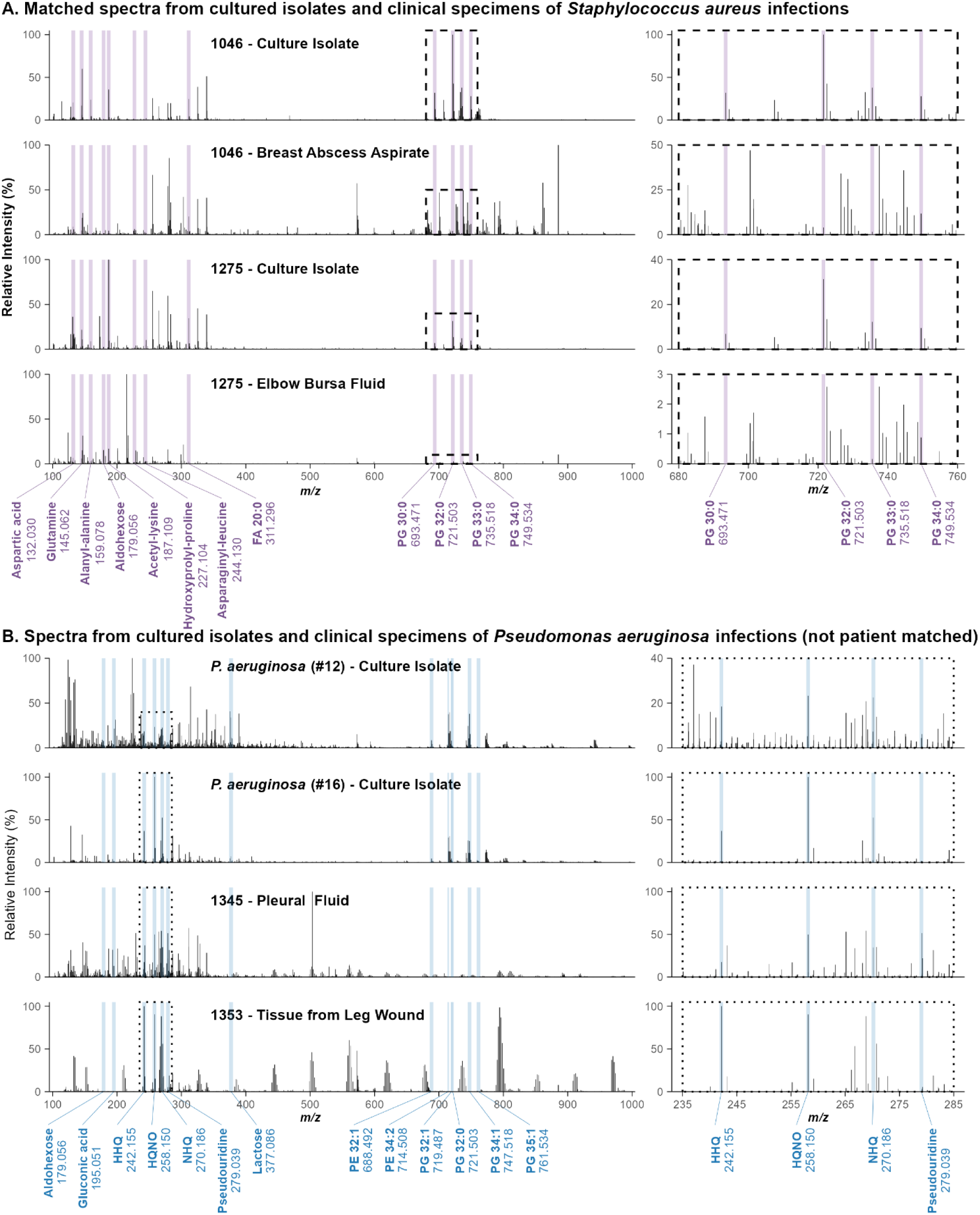
Comparison of MSPen spectra from clinical isolates and infected clinical specimens for full spectra. **(A)** MSPen spectra of *S. aureus* culture isolates and their corresponding infected clinical specimen. Sample code 1046 (top two spectra) corresponds to an infected breast abscess and sample code 1275 (bottom two spectra) corresponds to a right elbow bursa, both exhibited abundant bacterial growth in culture. Peaks highlighted in purple are those with positive Lasso feature weights for *S. aureus*. **(B)** MSPen spectra of *P. aeruginosa* culture isolates (top two spectra) and non-matched infected clinical specimens (bottom two spectra). Sample code 1329 corresponds to infected tissue and sample code 1345 corresponds to infected pleural fluid that had a thick, white appearance. Peaks highlighted in blue are those with positive Lasso feature weights for *P. aeruginosa*. Quorum-sensing molecules such as *m/z* 242.150 (2-heptyl-4-quinolone (HHQ) and *m/z* 258.150 (2-heptyl-4-hydroxyquinoline N-oxide (HQNO) are characteristic of *P. aeruginosa*. The right panels are magnified views of a portion of the lipid region between *m/*z 670-730 in (A) and (B) and the quinolone region between *m/*z 230-270 in (B).

We also analyzed 11 clinical specimens infected with a monomicrobial Gram-negative bacteria. Among them, 6 clinical specimens were infected with *P. aeruginosa*, including an aspirate, lung tissue, left leg ulcer, pleural fluid, and infected tissue samples. Unlike other bacteria, the MSPen mass spectra of *P. aeruginosa*-infected specimens presented characteristic *Pseudomonas* quinolone signal (PQS)-associated ions including *m/z* 242.155 (HHQ), *m/z* 258.150 (HQNO) and *m/z* 270.186 (NHQ) **(Fig. 3B)**. Representative MSPen mass spectra for infected pleural fluid and tissue (right leg wound), clinical specimens, and non-matched clinical isolates of *P. aeruginosa* are provided in **Fig. 3B** for comparison. Although the number and type of quinoline molecules detected was variable in the infected specimens analyzed, detection of even one of these species-specific molecules provides direct molecular evidence of *P. aeruginosa* infection (**Fig. S20**). In addition, rhamnolipids (Rha) including *m/z* 503.322 (Rha-C10-C10) and *m/z* 649.381 (Rha-Rha-C10-C10) were detected in pleural fluid and right leg wound tissue. Other clinical specimens infected with a monomicrobial Gram-negative bacteria included samples of prostate tissue abscess (*E. coli*), knee wound (*E. cloacae* complex), and cerebrospinal fluid (*K. pneumoniae*). Across clinical specimens infected with *Enterobacteriaceae* family bacteria (*K. pneumoniae*, *E. coli*, and *E. cloacae* complex), the mass spectra obtained exhibited characteristic lipids at *m/z* 702.508 (PE 33:1), *m/*z 719.487 (PG 32:1), *m/*z 733.503 (PG 33:1), *m/*z 745.503 (PG 34:2), *m/z* 747.518 (PG 34:1), and *m/*z 773.534 (PG 36:2) (**Fig. S21 and S22**). These molecular features matched those observed in isolate mass spectra and served as consistent markers of Gram-negative infection in clinical specimens analyzed.

The remaining 3 infected clinical specimens presented polymicrobial infections, in which two samples contain both Gram-positive and Gram-negative bacteria and yielded MSPen mass spectra with mixed molecular signatures characteristic of both groups. For example, an infected right leg wound sample (sample 1281) showed heavy growth of *K. pneumoniae*, *Klebsiella oxytoca*, and *Proteus mirabilis*, along with moderate growth of *S. aureus* and *E. faecalis*. Gram-negative associated lipid ions at *m/z* 702.508 (PE 33:1), *m/z* 719.487 (PG 32:1), and *m/*z 733.503 (PG 33:1) were detected alongside Gram-positive associated peaks at *m/z* 735.518 (PG 33:0) and 749.534 (PG 34:0), reflecting the coexistence of multiple bacterial taxa within the same specimen (**Fig. S23**). In a representative polymicrobial specimen from a left posterior thigh wound (sample 1269) containing *P. aeruginosa* (4+), *P. mirabilis* (4+), *K. oxytoca* (2+), *Streptococcus oralis* (2+), *and Klebsiella variicola* (1+), MSPen mass spectra revealed PQS molecules at *m/z* 242.155 (HHQ), and *m/z* 270.186 (NHQ) in addition to lipid peaks at *m/z* 735.518 (PG 33:0), *m/z* 747.518 (PG 34:1), and *m/z* 749.534 (PG 34:0), providing evidence of both Gram-negative and Gram-positive bacterial populations within the same infected specimen (**Fig. S20**). The third clinical specimen, left knee bursa tissue (sample 1221), exhibited heavy growth (4+) of *S. pyogenes* and low growth (1+) of *S. aureus*. Both bacteria belong to Gram-positive genera, and we expected to observe characteristic lipids from the *Streptococcus* genus due to heavier growth from *S. pyogenes*. The characteristic features observed at *m/z* 735.518 (PG 33:0), *m/z* 745.503 (PG 34:2), *m/z* 747.518 (PG 34:1), *m/z* 771.518 (PG 36:3), *m/z* 773.534 (PG 36:2), and *m/z* 775.550 (PG 36:1) matched those identified in the clinical isolate of *S. pyogenes* (**Fig. S24**).

To benchmark MSPen-based classification against MALDI-TOF, five of the infected clinical specimens analyzed were randomly selected (sample 1063: prosthetic right hip joint fluid, *S. epidermidis*; sample 1275: right elbow bursa fluid, MSSA; sample 1277: left hip fluid, MSSA; sample 1359: thoracic fluid, MRSA; and sample 1046: breast abscess, MSSA), cultured on blood agar, and the clinical isolates were analyzed by MALDI-TOF and MSPen. MALDI-TOF correctly identified all samples as *Staphylococcus* species (four *S. aureus*, one *S. epidermidis*). Similarly, statistical classification using the Gram and species relative intensity lasso models correctly identified all isolates derived from the clinical specimens as Gram-positive, with four classified as *S. aureus* and one as *S. epidermidis*, in full agreement with the MALDI-TOF results **(Table S1)**.

### Disturbance in lipid isotopic ratio patterns in infected clinical specimens

We next investigated whether bacterial infection in human specimens alters the natural ratios of relative abundances of host human lipids. We first evaluated the lipid ion at *m/z* 747.518 (PG 34:1), which was consistently observed in both Gram-positive and Gram-negative bacteria (**Fig. 1B**) and commonly detected as a naturally occurring lipid in human tissues with the MSPen and other MS techniques (20, 26). The MSPen mass spectra of the infected clinical human specimens indicated that the ratio of this bacterial lipid compared to lipids that are exclusively detected in host human tissues appeared elevated in infected human specimens due to the shared source of this lipid. To validate this observation, we performed MSPen analysis of 67 non-infected human breast and muscle tissues (n = 10 breast, n = 23 skeletal muscle, and n = 34 smooth muscle) and calculated the lipid ratio of *m/z* 747.518 (detected in both host and bacteria isolates) to *m/z* 748.528 (PE O-38:6; only detected in human tissues) in the 57 sterile tissues and the 39 infected human specimens. Consistent with our observation, the log-ratio of *m/z* 747.518 to *m/z* 748.528 was significantly higher in infected clinical specimens (ratio = -0.92) compared with non-infected muscle and breast tissue (ratio = -1.59) (**Fig. 4A**; Wilcoxon rank-sum test, p = 4.4 x 10⁻^5^). This result suggests enrichment of shared bacterial-human lipid signals relative to host-only lipid signals in infected patient specimens.

**Figure 4.**
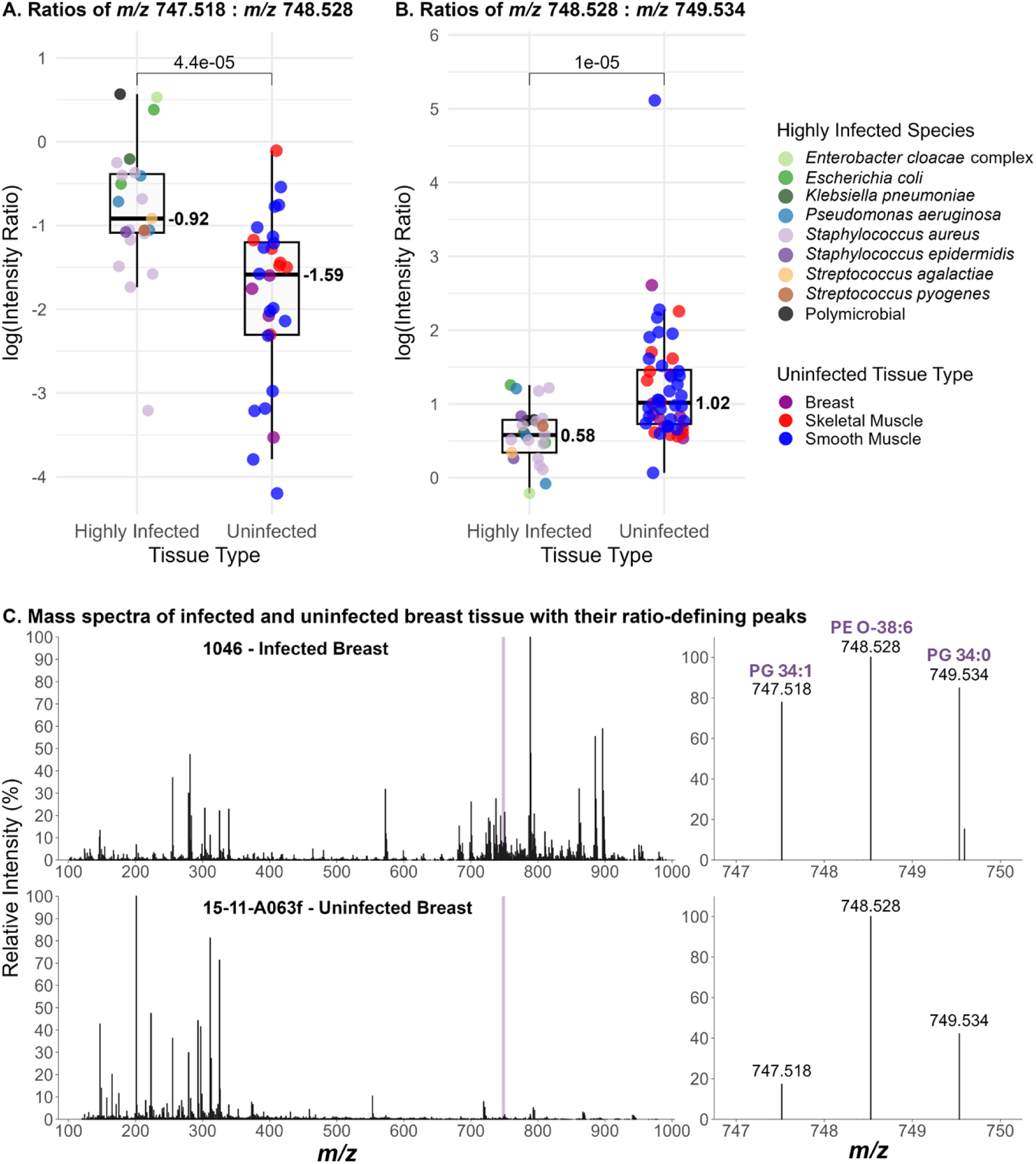
Box plots showing the log-intensity ratios of key lipid ions in infected and non-infected tissues. **(A)** log-intensity ratio of *m/z* 747.518 : *m/z* 748.528; **(B)** log-intensity ratio of *m/z* 748.528 : *m/z* 749.534. **(C)** Mass spectra of an infected breast abscess (top) and non-infected breast tissue (bottom), with an expanded view of the *m/z* 747–750 region shown on the left. The peak at *m*/*z* 747.518, identified as PG 34:1, is commonly detected in *Staphylococcus* and *Streptococcus* species, as well as in some Gram-negative species like *P. aeruginosa*, but is relatively lower in human tissue. The peak at *m/z* 749.534, identified as PG 34:0, is primarily associated with Gram-positive *Staphylococcus* species, especially *S. aureus* and *S. epidermidis*. The ion at *m/z* 748.528, assigned as PE O-38:6, is observed exclusively in human tissue and not in any bacterial isolates. Therefore, in infected samples, the relative intensity of *m/z* 747.518 is expected to increase due to the additional contribution from bacterial lipid signals. In Figure 4A, as a result, the log-ratio (*m/z* 747.518: *m/z* 748.528) becomes higher in infected tissues compared with non-infected muscle. In B, the log ratio of *m/z* 748.528: *m/z* 749.534 is relatively lower in infected tissues. This is partly due to the overlap between *m/z* 749.534 (PG 34:0) and the second isotope of *m/z* 748.528, which elevates the apparent intensity at *m/z* 749.534 in infected samples. In Panel 4C (left), a clear difference in the isotopic pattern is observed between the infected breast abscess and the uninfected breast tissue. Together, these findings highlight a promising lipid ion ratio-based approach for distinguishing infected from non-infected tissues using MSPen.

We then examined whether lipids from bacteria also perturb the natural isotopic distribution of host-derived lipids in the 39 infected patient specimens analyzed. In the uninfected human tissues, the ion at *m/z* 748.528 (only detected in human tissues) and its ^13^C isotopic peak at *m/z* 749.534 have an expected naturally occurring intensity ratio of ∼100:48. In contrast, in highly infected specimens, this isotopic ratio was markedly altered to ∼100:90. Accordingly, the log-ratio of *m/z* 748.528 to *m/z* 749.534 was significantly lower in infected specimens (ratio = 0.58) than in non-infected muscle and breast tissues (ratio = 1.02) (**Fig. 4B**, Wilcoxon rank-sum test, p = 1.0 × 10⁻^5^). This distortion arises from the overlap between host ^13^C lipid isotopic signal of PE O-38:6 and the ^12^C bacterial lipid signal at the same nominal *m/z* value of 749.534. In *S. aureus*, the ion at *m/z* 749.534 corresponds to a bacterial lipid PG 34:0 (molecular formula C_40_H_79_O_10_P; ionic form [M-H]^-^; mass accuracy ±0.005 Da). As bacterial burden increases, the bacterial PG 34:0 signal at *m/z* 749.534 overlaps with and dominates the host PE O-38:6 ^13^C isotope peak, resulting in a decrease of the *m/z* 748.528/749.534 ratio from 2:1 to approximately 1:1 (**Fig. 4B and 4C**). This effect indicates dominance of bacterial lipid contribution at *m/z* 749.534 in infected tissue specimens when compared to sterile human tissues. Disturbed isotopic ratios between *m/z* 748.528 and *m/z* 749.534 were consistently observed across infected clinical samples.

## Discussion

To address the challenge of bacterial identification in clinical samples, we employed the MSPen technology for direct, non-destructive, and rapid molecular analysis of clinical isolates and infected human specimens. We analyzed 755 pure clinical isolates representing the most frequently occurring bacterial pathogens, including *E. faecium, S. aureus, K. pneumoniae, P. aeruginosa, and* the *Enterobacte*r species. The mass spectral data obtained was characterized by metabolites and lipids previously detected in bacterial isolates with MS techniques. From the data obtained, we generated a comprehensive molecular database that enabled construction of a hierarchical classification framework encompassing Gram-positive versus Gram-negative discrimination, *Staphylococcus* versus *Streptococcus* genus-level classification, GAS vs. GBS differentiation, and species-level identification of pure isolates.

For Gram typing, the classification models generated with MSPen data achieved greater than 98% accuracy for Gram-type identification across training and test sets. Similar performance was achieved for *Staphylococcus* and *Streptococcus* genera classification (>98% accuracy), and GAS versus GBS classification (>90% accuracy with relative intensity lasso and 100% accuracy with log-ratio lasso). At the species level, classification using multinomial logistic regression with lasso, log-ratio lasso, and random forest achieved high overall accuracies ranging from 92-96%. Misclassifications were primarily observed among species with highly similar lipid profiles in the *m/z* 650-850 range, particularly between *S. aureus* and *S. epidermidis*, as well as among *Enterobacteriaceae* species (*E. coli*, *E. cloacae* complex and *K. pneumoniae*). These results indicate that while intrinsic biochemical similarity in metabolic species is a potentially limiting factor for MSPen classification of closely related species, profiles of metabolites and lipids and the ratios of their relative abundances provide a powerful means for bacterial identification from isolates. Notably, these results are comparable to current metrics of clinical-based molecular tests for bacterial identification. In the case of MALDI-TOF, microbial identification of cultured isolates is based on the detection of proteins and non-ribosomal peptides in the mass range of ∼2,000-15,000 Da. The mass spectrum is matched to reference libraries built from thousands of bacteria-specific protein patterns, yielding an identification confidence score for a specific genus or species. Commercial MALDI-TOF systems have reported accuracies ranging from 90-95% at the genus and species levels, which are comparable to the accuracies achieved in our study with classifiers built with a much smaller dataset of metabolites and lipids MSPen data. Although bacterial proteins are generally considered to be more specific than metabolites and lipids, closely related bacterial species also share protein markers that can also confound MALDI-TOF identification. Higher specificity for bacteria identification is achieved with 16S rRNA typing of bacterial RNA, with reported accuracies in the range of ∼97-100%. However, 16S rRNA requires careful primer design and is more time consuming than MALDI-TOF. Complementary to both MALDI-TOF and 16 sRNA, MSPen analysis is performed in seconds directly on the clinical isolate and without any sample preparation, thus providing an alternative approach with similar performance that could be valuable in clinical practice for isolates.

Among the lipid species detected by MSPen and selected as predictive markers in our classifier, *m/z* 693.472 (PG 30:0), *m/z* 721.503 (PG 32:0), *m/z* 735.518 (PG 33:0), and *m/z* 749.534 (PG 34:0) were found at relatively higher abundance in *S. aureus*. We observed a similar lipid peak pattern in our previous MSPen-based bacterial identification study (*29*). The consistent trend observed across both the previous and current studies highlight the reproducibility of the phosphatidylglycerol (PG) lipid species in *S. aureus*. Therefore, this lipid class appears to represent an important biomarker group for infection studies and bacterial discrimination. Amongst all organisms we analyzed, *P. aeruginosa* exhibited the most distinctive molecular signature characterized by prominent peaks consistently detected within the *m/z* 200-350 range. These ions were identified as quorum-sensing molecules belonging to the 2-alkylquinolone family, which regulates population density and virulent gene expression in *P. aeruginosa* (*30–32*). Key PQS molecules were detected at *m/z* 242.155 (HHQ), *m/z* 258.150 (HQNO), and *m/z* 270.186 (NHQ), along with mono- and di-rhamnolipids, providing highly specific molecular markers for this pathogen. These findings underscore the value of small molecule-signaling metabolites as diagnostic markers when lipid profiles alone are insufficiently specific.

We also evaluated whether the metabolic and lipid profiles detected by the MSPen would enable discrimination between MRSA and MSSA, a difficult but critical task that has major clinical implications in selection of antibiotic treatment. The prediction accuracy achieved with classification methods ranged from 67–69% in the training and test sets for logistic regression (overall accuracy), 62% in both the training and test sets for log-ratio lasso (overall accuracy), and 65% overall accuracy in the test set for the random forest model. The moderate performance results are coherent with the fact that the mass spectra data obtained showed similar molecular profiles without any major differences in abundance of specific ions or molecular species (**Fig. S15**). Although previous studies have shown distinct patterns of lipids between MRSA and MSSA with LC-MS (*33*), progress in direct MS analysis of lipids for MRSA and MSSA discrimination has been limited. Previous work with MALDI-MS imaging using over 20,000 clinical isolates has shown greater promise for MSSA and MRSA discrimination based on peptide and protein analysis (*34*), yielding performance with an area under the receiver operating characteristic curve of 0.78 to 0.88. Based on the large number of isolates used by MALDI, we plan to pursue further work employing MSPen towards a much larger sample size of MRSA and MSSA strains for improved statistical power and to optimize assays to deepen the range of molecular analyses.

To assess whether metabolites and lipid features detected by the MSPen in pure isolates could be observed in infected clinical samples, we performed MSPen analysis of a diverse set of highly infected human specimens prospectively collected for our study, including abscesses, tissues, aspirates, cerebrospinal fluid, and synovial fluid. To identify molecular changes that were specific to infections, we also analyzed sterile human muscle and breast tissues as control tissues. In clinical specimens infected with *Enterobacteriaceae* species *(E. coli*, *K. pneumoniae*, and *E. cloacae* complex), for example, characteristic lipids observed in *Enterobacteriaceae* isolates, including *m/z* 702.508 (PE 33:1), *m/z* 733.503 (PG 33:1), and *m/*z 747.518 (PG 34:1), were detected (**Fig. S21, S22**). In specimens infected with *S. aureus*, which were majority biological fluids including bursa fluid, prosthetic joint fluid, synovial fluid, and breast abscess, characteristic lipid peaks at *m/z* 735.518 (PG 33:0) and 749.534 (PG 34:0) were detected (**Fig. S25**). In these samples, the intensity of the *m/z* 749.534 peak was markedly elevated relative to uninfected human tissues, resulting in a disturbance of the expected natural isotopic ratio of the adjacent host lipid at *m/z* 748.528 (PE O-38:6). The observation that these lipid peaks are detected in infected tissues are consistent with prior MALDI imaging studies. For example, Perry *et al*. reported detection of PG species at *m/*z 693.471 (PG 30:0), *m/*z 707.481 (PG 31:0), *m/*z 721.502 (PG 32:0), and *m/*z 735.518 (PG 33:0) in cardiac and kidney tissues infected with *S. aureus*(*35*), while Good *et al.* observed similar PG lipids in infected bone tissue (*36*). Our findings extend these observations by demonstrating direct detection of these bacterial lipids from intact clinical specimens using the MSPen and the effect of overlapping lipid species in isotopic distributions in all infected human specimens analyzed. In the unique case of specimens infected with *P. aeruginosa*, detection of PQS molecules directly from clinical samples provided a reliable and specific means of bacteria identification even when host and bacterial lipids overlapped. These results are supported by prior reports of PQS detection in sputum from cystic fibrosis patients infected with *P. aeruginosa* using LC-MS/MS, reinforcing the translational relevance of quorum-sensing molecules as infection biomarkers (*37*).

It is important to note that our study has limitations. First, the MSPen mass spectra profiles acquired from isolates are dependent on the culture medium of growth, which must be considered when evaluating molecular profiles and building statistical classifiers. For example, *m/z* 731.487 (PG 33:2), *m/z* 733.502 (PG 33:1), and *m/z* 735.518 (PG 33:0) were clearly detected in blood agar-grown *S. aureus*. In contrast, PG 33:2 and PG 33:1 showed lower intensity in nutrient agar–grown cultures, indicating medium-dependent differences in lipid expression (**Fig. S26**). In this study, isolates were cultured on blood agar, nutrient agar, or chocolate agar as appropriate (*S. aureus* (n = 328; n = 48 on nutrient agar and n = 280 on blood agar); *E. coli, P. aeruginosa, K. pneumoniae, and Streptococcus spp.* on blood agar; and *H. influenzae* on chocolate agar), which allowed direct comparison of molecular trends among the 755 isolates analyzed. Another challenge in our study is related to the ability to predict bacteria identification from the MSPen mass spectra obtained from infected clinical specimens. Although many of the bacterial molecules detected from isolates were detected directly from infected clinical samples with the MSPen, differences in the mass spectra acquired from isolates and clinical samples were observed. These differences, as expected, primarily arise from the background of biological peaks from the human-host cells, as well as potential infection site chemistry, and other exogenous matrix compounds. Infected tissues are cellularly dense and, inevitably, the MSPen mass spectra obtained presented high relative intensity of host-lipids that suppress detection of bacterial molecules. For example, while *m/z* 721.503 (PG 32:0) was detected at highest relative abundance in the MSPen mass spectra of *S. aureus* and *S. epidermidis* isolates (**Figure 1B**), this ion was detected in much lower relative abundance in infected breast tissue tissues, in which host-cellular lipids dominate the mass spectra (**Figure 3A**). In infected fluids, salt cluster ions were frequently detected in the mass spectra, often suppressing the detection of bacterial molecules (**Fig. S27**). Salt cluster ions were also detected in tissues that had been exposed to saline during sample processing, which will be avoided in future studies. Lastly, we only focused our study on clinical specimens with high bacterial burden (3+ or 4+) to maximize the sensitivity for detection of bacterial molecules within the host-molecular background. While species-level classification in infected specimens remains challenging, the detection of pathogen-specific molecular signatures provides molecular confirmation of infection.

Overall, our study demonstrates that MSPen-based molecular profiling enables rapid (∼10 seconds), direct detection of bacterial metabolic signatures from clinical isolates, with promising performance for the detection of infection directly from clinical specimens. Future studies are focused on expanding and optimizing our MSPen analysis in the positive ion mode to capture known lipids and peptides that are unique to bacteria, such as species-specific Lipid A/peptide toxins (*38–40*). Continuous development and refinement of our statistical classifiers are also being pursued to increase predictive performance by including larger and more complex sample sets, including larger sample sets of MRSA and MSSA strains. Analyses of other clinically relevant microbial species including fungus and polymicrobial infections are also being pursued to expand the value of this technology in clinical microbiology. Lastly, advanced analytical and MS methods that increase sensitivity for detection of bacterial molecules from samples with lower bacterial bunder (<3+) are being explored to allow detection of infection in human samples with different bacterial loads. Collectively, our study shows that by eliminating the need for extensive sample preparation and culturing, this approach has the potential to support real-time bacterial detection in clinical and intraoperative settings, facilitating faster therapeutic decision making and improved patient care.

## Materials and Methods

### Clinical Samples

A total of 794 (755 pure isolates and 39 infected specimens) clinical samples were obtained from Dell Children’s Medical Center in Austin, Texas, Riley Hospital for Children in Indianapolis, Indiana, Baylor College of Medicine in Houston, Texas, and Texas Children’s Hospital in Houston, Texas under approved IRB protocol (for TCH IRB H-51585). A total of 755 pure isolates were analyzed, including *Enterobacter cloacae complex* (n = 40), *E. coli* (n = 45), *H. influenzae* (n = 41), *K. pneumoniae* (n = 50), *P. aeruginosa* (n = 81), *E. faecalis* (n = 13), *E. faecium* (n = 14), *S. epidermidis* (n = 48), *S. aureus* (including MRSA, n = 158, and MSSA, n = 170), *S. pyogenes* (GAS, n = 40), *S. agalactiae* (GBS, n = 27), and *S. pneumoniae* (n = 28). Clinical specimens (n = 46) were surgically collected from patients as part of routine care (**Supplementary Data Excel Sheet 5**). Further analysis was performed on a subset of 36 samples infected with only one microbial species, and whose infection criteria consisted of 3+ (moderate) and 4+ (abundant) growth in standard culture media. Banked uninfected normal muscle and breast tissue samples were obtained from the Cooperative Human Tissue Network (CHTN) and MD Anderson Cancer Center (MDACC). Isolated bacterial colonies were recovered on 5% sheep blood or chocolate agar plates (Remel) after overnight incubation at 35°C in 5% CO2. Identification was confirmed using conventional clinical laboratory methods such as MALDI-TOF mass spectrometry (Vitek MS, bioMerieux) or biochemical panels (Vitek 2, bioMérieux). A desalting step was performed for samples stored in normal saline to reduce salt interference during MS analysis (**Supplemental Methods**).

### MSPen Analysis

The MSPen consists of a handheld sampling probe, a microcontroller unit, and a mass spectrometer. Analyses were conducted with a MSPen tip diameter of 5 or 7 mm. 100% methanol was used as the solvent which is delivered at 800 µL/min for 2 seconds. The solvent droplet deposited by the MSPen desorbed molecules from the sample (extraction time) for 7 seconds. The tubing length between the MSPen tip and mass spectrometer was 0.5 m. The total analysis time from initiating the MSPen controller to deposit a solvent droplet to analysis by the mass spectrometer was 15 seconds. The analyses were conducted with a ThermoFisher Scientific Exploris 120 Orbitrap mass spectrometer, which has a resolving power of 120,000. The inlet capillary temperature was set to 325°C and the ion source temperature was set to 350°C. Mass spectra were collected in negative ion mode and with a detection range of *m/z* 100-2000.

All analysis procedures followed the guidelines outlined in the Institutional Biosafety Committee protocol. For MSPen analysis of agar cultures, colonies were removed from the agar plate with a sterile plastic inoculating loop and smeared onto a glass slide which were then directly analyzed with the MSPen. The bacterial isolates were examined in different batches using the MSPen in a randomized order. Clinical specimens were analyzed directly with the MSPen or desalted and analyzed as described above.

Metabolite and lipid ion peaks were tentatively identified using LIPID MAPS, the Metabolomics Workbench, the Human Metabolome Database, the *E. coli* Metabolome Database (ECMDB), and the Dictionary of Natural Products, generating a library of over 200 molecules in the *m/z* 100-1000 range (**Supplementary Data Excel Sheet 1**)(*41–45*).

### Statistical analysis and Data Visualization

Raw MSPen data were extracted to CSV format in centroid mode using ThermoFisher XCalibur 3.0.67 and FreeStyle™ 1.8 SP2, Modern Data Visualization Software, Version 1.8.63.0. Further data preprocessing and statistical analysis was performed in R (version 4.5.3). Total ion count (TIC) normalization was performed on all samples. Samples were labeled with their respective class for the desired analysis (gram typing, species, or sub-species specifications). Pure cultured samples were randomly split at the patient level into 70% for training and 30% for test, while all clinical isolates were used as a testing set. Relative intensity lasso with both logistic and multinomial regression were performed using the ‘glmnet’ package (version 4.1-8) in the R CRAN language library (*46*). Training set performance was evaluated using leave-one-out CV. Accuracy, sensitivity, and specificity were calculated against known class labels. Random forests were performed using the ‘ranger’ (version 0.18.0) and ‘caret’ (version 7.0-1) packages (*47, 48*). Near zero variance peakswere removed using the “nearZeroVar” function from ‘caret’, which excludes features with either zero variance or low variability that meet both of the following criteria: (i) a high frequency ratio (>19) between the most and second most common values, and (ii) a low proportion of unique values (<10%) across samples. Repeated 5X CV with five repeats was performed using a tuning grid with the “gini” and “extratrees” split rules. Test set performance was evaluated similarly to lasso models.

As noted, log-ratio lasso analysis was also performed. After data preprocessing and normalization, pairwise logarithmic ratio calculations (x) were performed across all selected *m/z* peaks in which MS/MS was performed, so that the number of new ratio features (matrix columns) was equal to 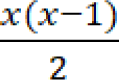 and so that the data input now contained ratios of normalized intensity values in the form natural 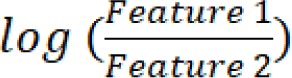 . In cases where one or both input feature intensities were 0, the log-ratio was calculated as positive or negative infinity, or an undefined value. The lasso analysis performed in ‘glmnet’ cannot effectively manage these infinite or undefined values and removing them from analysis would result in very few finite features. Thus, positive infinity values were imputed to the 99th percentile of each matrix column’s log-ratio intensity values, negative infinity to the 1st percentile, and undefined values toward zero. This matrix of log-ratio values was then used as the input for lasso as described previously. Ratio plots were created by performing natural log-ratio calculations for selected features between highly infected clinical samples and normal muscle without data imputation. Plots were created using the ‘ggplot2’ (version 4.0.0) package in R(*49*). Statistical analysis between the calculated finite log-ratios for all highly infected clinical samples and normal muscle was performed by Wilcoxon rank-sum test. Infinite and undefined log-ratio values were not included during statistical analysis or for plotting, thus only ratios resulting from samples with non-zero relative abundance at the specified peaks were included.

To visualize the cultured isolates, dimensionality reduction was performed after data preprocessing and normalization. PCA was performed on the entire dataset and all PCs were used as the t-SNE initialization. The variance coverage across the first 36 PCs are provided in **Fig. S28**. t-SNE was performed using the ’Rtsne’ (version 0.17) package in R(*50–52*), with the following parameters: perplexity = 15, max_iter = 7000, pca = FALSE, and normalize = FALSE. These decisions were selected based on running multiple versions with a range of perplexity values and the use of preprocessing, normalization, and PCA prior to t-SNE analysis.

## Data and Materials Availability

The data for this study are available for researchers and have been deposited.

## Supporting information

Pdf and Excelsheets

## Acknowledgements.

This work was funded by the NIH under grant R21HD106614. Additionally, Dr. Kirkpatrick receives salary support and research funding under K12HD111057 and P30HD106451. We thank Baylor College of Medicine, Department of Surgery, Houston, TX, USA; Texas Children’s Hospital, Department of Pathology, Houston, TX, USA; Dell Children’s Medical Center, Austin, TX, USA; and Riley Hospital for Children – IU Health, Department of Pediatric Infectious Diseases, Indianapolis, IN, USA, for providing clinical isolates and infected samples. We thank Mingxun Wang (UC Riverside) for insightful discussions.

## Author contributions

M.K., L.M.K., and L.S.E. designed research; M.K., F.E.J., J.I.M., and L.S.E. performed research; M.K., S.B., and A.E.M. performed tissue analysis; M.K., F.E.J., J.I.M., W.B., and J.H. provided statistical analysis and analytical tools; C.L.J, J.J.D., L.M.K., S.B.M, and R.D.D. contributed reagents and isolates; M.K., F.E.J., J.I.M., and L.S.E. wrote the paper. All authors reviewed and approved the paper prior to submission.

## Supplementary Materials

The PDF file includes:

Supplementary Text

Figs. S1 to S28

Table S1

Excel Sheets 1 to 5

