## Supplementary material for "Rapid Identification of Infections Directly from Isolates and Clinical Specimens with the MasSpec Pen Technology": Pdf and Excelsheets: SI_MSPen Bacteria-6_15_2026.pdf

A)

| Training Set | | Predict | |
|---|---|---|---|
| | Gram | Gram Negative | Gram Positive |
| True | Gram Negative | 174 | 6 |
| | Gram Positive | 3 | 342 |

| Class | Recall Rate |
|---|---|
| Gram | 96.7% |
| Gram Positive: | 99.1% |
| Overall | 98.3% |

B)

| Test Set | | Predict | |
|---|---|---|---|
| | Gram | Gram Negative | Gram Positive |
| True | Gram Negative | 77 | 0 |
| | Gram Positive | 0 | 153 |

| Class | Recall Rate |
|---|---|
| Gram | 100.0% |
| Gram Positive: | 100.0% |
| Overall | 100.0% |

**Figure S1.** Logistic regression lasso for Gram stain classification of pure clinical isolates. **(A)** Confusion matrix of the training set and **(B)** test set.

A)

| Training Set | | Predict | |
|---|---|---|---|
| | Gram | Gram Negative | Gram Positive |
| True | Gram Negative | 175 | 5 |
| | Gram Positive | 6 | 339 |

| Class | Recall Rate |
|---|---|
| Gram | 97.2% |
| Gram Positive: | 98.3% |
| Overall | 97.9% |

B)

| Test Set | | Predict | |
|---|---|---|---|
| | Gram | Gram Negative | Gram Positive |
| True | Gram Negative | 77 | 0 |
| | Gram Positive | 1 | 152 |

| Class | Recall Rate |
|---|---|
| Gram | 100.0% |
| Gram Positive: | 99.3% |
| Overall | 99.6% |

**Figure S2.** Log-ratio lasso for Gram stain classification of the pure clinical isolate **(A)** training and **(B)** test sets.

| Test Set | | Predict | |
|---|---|---|---|
| | Gram | Gram Negative | Gram Positive |
| True | Gram Negative | 77 | 0 |
| | Gram Positive | 1 | 152 |

| Class | Recall Rate |
|---|---|
| Gram | 100.0% |
| Gram Positive: | 99.3% |
| Overall | 99.6% |

**Figure S3.** Random forest for Gram stain classification of the pure clinical isolate test set.

A)

| Training Set | | Predict | |
|---|---|---|---|
| | Genus | *Staphylococcus* | *Streptococcus* |
| True | *Staphylococcus* | 259 | 0 |
| | *Streptococcus* | 1 | 66 |

| Class | Recall Rate |
|---|---|
| *Staphylococcus*: | 100.0% |
| *Streptococcus*: | 98.5% |
| Overall | 99.7% |

B)

| Test Set | | Predict | |
|---|---|---|---|
| | Genus | *Staphylococcus* | *Streptococcus* |
| True | *Staphylococcus* | 116 | 1 |
| | *Streptococcus* | 0 | 28 |

| Class | Recall Rate |
|---|---|
| *Staphylococcus*: | 99.1% |
| *Streptococcus*: | 100.0% |
| Overall | 98.6% |

**Figure S4.** Logistic regression lasso discrimination of *Staphylococcus* versus *Streptococcus* **(A)** training set and **(B)** test set.

A)

| Training Set | | Predict | |
|---|---|---|---|
| | Genus | *Staphylococcus* | *Streptococcus* |
| True | *Staphylococcus* | 255 | 4 |
| | *Streptococcus* | 1 | 66 |

| Class | Recall Rate |
|---|---|
| *Staphylococcus*: | 98.5% |
| *Streptococcus*: | 98.5% |
| Overall | 98.5% |

B)

| Test Set | | Predict | |
|---|---|---|---|
| | Genus | *Staphylococcus* | *Streptococcus* |
| True | *Staphylococcus* | 117 | 0 |
| | *Streptococcus* | 0 | 28 |

| Class | Recall Rate |
|---|---|
| *Staphylococcus*: | 100.0% |
| *Streptococcus*: | 100.0% |
| Overall | 100.0% |

**Figure S5.** Log-ratio lasso discrimination of *Staphylococcus* versus *Streptococcus* **(A)** training and **(B)** test set.

| Test Set | | Predict | |
|---|---|---|---|
| | Genus | *Staphylococcus* | *Streptococcus* |
| True | *Staphylococcus* | 117 | 0 |
| | *Streptococcus* | 0 | 28 |

| Class | Recall Rate |
|---|---|
| *Staphylococcus*: | 100.0% |
| *Streptococcus*: | 100.0% |
| Overall | 100.0% |

**Figure S6.** Random forest classification of *Staphylococcus* vs *Streptococcus* on the test set.

A)

| Training Set | | Predict | |
|---|---|---|---|
| | Species | *GAS* | *GBS* |
| True | *GAS* | 28 | 0 |
| | *GBS* | 1 | 18 |

| Class | Recall Rate |
|---|---|
| GAS: | 100.0% |
| GBS: | 94.7% |
| Overall | 97.9% |

B)

| Test Set | | Predict | |
|---|---|---|---|
| | Group | GAS | GBS |
| True | GAS | 12 | 0 |
| | GBS | 2 | 6 |

| Class | Recall Rate |
|---|---|
| GAS: | 100.0% |
| GBS: | 75.0% |
| Overall | 90.0% |

**Figure S7.** Logistic regression lasso discrimination of GAS (*S. pyogenes*) versus GBS (*S. agalactiae*) on the **(A)** training set and **(B)** test set.

A)

| Training Set | | Predict | |
|---|---|---|---|
| | Species | *GAS* | *GBS* |
| True | *GAS* | 28 | 0 |
| | *GBS* | 0 | 19 |

| Class | Recall Rate |
|---|---|
| GAS: | 100.0% |
| GBS: | 100.0% |
| Overall | 100.0% |

B)

| Test Set | | Predict | |
|---|---|---|---|
| | Group | GAS | GBS |
| True | GAS | 12 | 0 |
| | GBS | 0 | 8 |

| Class | Recall Rate |
|---|---|
| GAS: | 100.0% |
| GBS: | 100.0% |
| Overall | 100.0% |

**Figure S8.** Log-ratio lasso discrimination of GAS (*S. pyogenes*) versus GBS (*S. agalactiae*) on the **(A)** training set and **(B)** test set.

| Test Set | | Predict | |
|---|---|---|---|
| | Group | GAS | GBS |
| True | GAS | 12 | 0 |
| | GBS | 0 | 8 |

| Class | Recall Rate |
|---|---|
| GAS: | 100.0% |
| GBS: | 100.0% |
| Overall | 100.0% |

**Figure S9.** Random forest classification of GAS (*S. pyogenes*) versus GBS (*S. agalactiae*) on the test set.

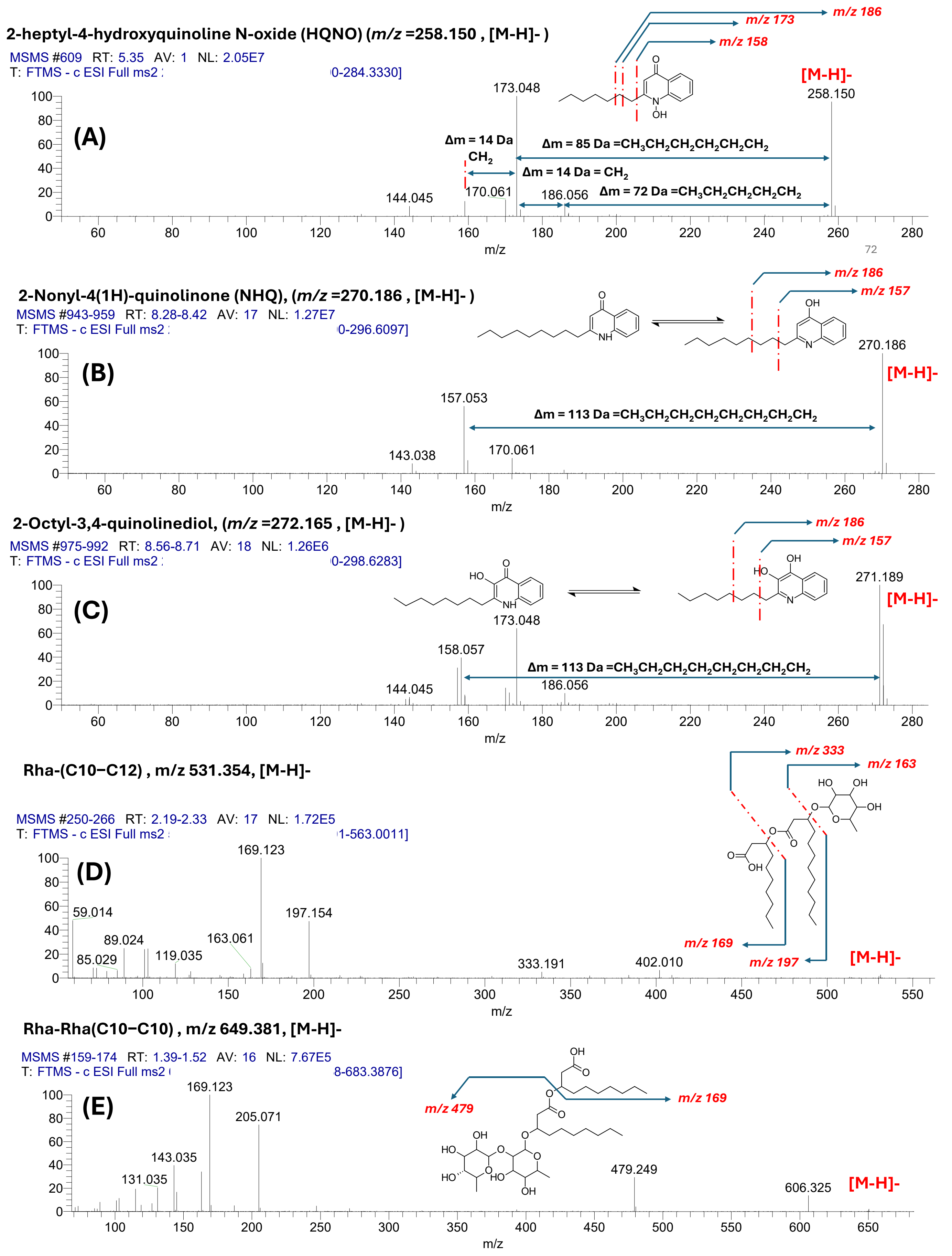

**Figure S10.** Tandem mass spectra (MS/MS) of selected PQS-class molecules and rhamnolipids: (A) *m/z* 258.150 (2-heptyl-4-hydroxyquinoline N-oxide [HQNO]); (B) *m/z* 270.186 (2-Nonyl-4(1H)-quinolinone [NHQ]), (C) *m/z* 272.165 (2-octyl-3,4-dihydroxyquinoline); (D) *m/z* 531.354 (mono-rhamnolipid [Rha–C22]); and (E) *m/z* 649.381 (di-rhamnolipid [Rha–Rha–C20]).

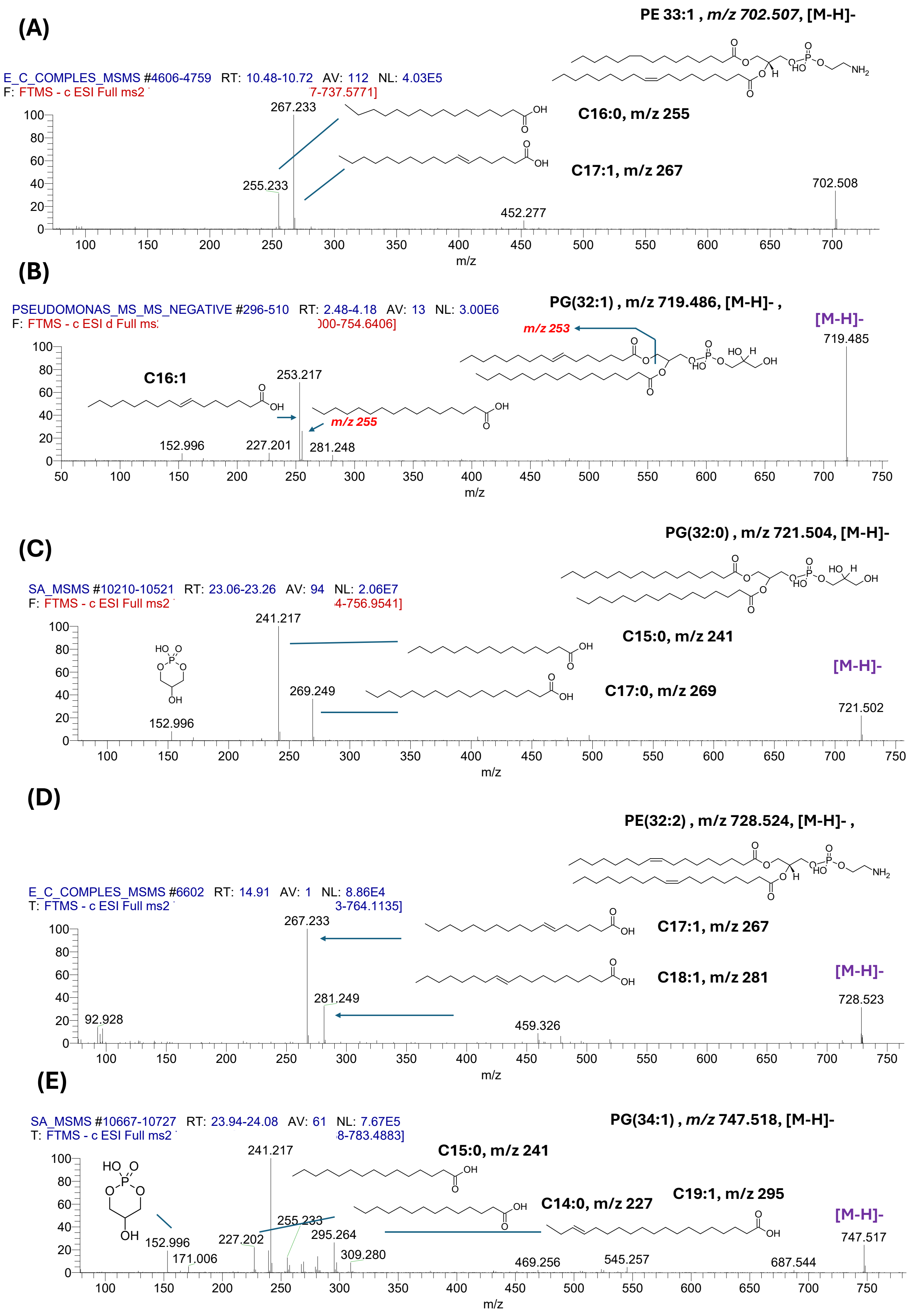

**Figure S11.** Tandem mass spectra (MS/MS) of selected phosphatidylethanolamine (PE) and phosphatidylglycerol (PG) lipid species: (A) *m/z* 702.507 (PE 33:1); (B) *m/z* 719.486 (PG 32:1); (C) *m/z* 721.504 (PG 32:0); (D) *m/z* 728.523 (PE 35:2); and (E) *m/z* 747.518 (PG 34:1).

**Training Set Confusion Matrix**

| | | Predict | | | | | | | | | | | |
|---|---|---|---|---|---|---|---|---|---|---|---|---|---|
| | **Species** | ***Enterobactercloacae complex*** | ***Enterococcus faecalis*** | ***Enterococcus faecium*** | ***Escherichia coli*** | ***Haemophilus influenza*** | ***Klebsiellapneumoniae*** | ***Pseudomonas aeruginosa*** | ***Staphylococcus aureus*** | ***Staphylococcus epidermidis*** | ***Streptococcus agalactiae*** | ***Streptococcus pneumoniae*** | ***Streptococcus pyogenes*** |
| True | ***Enterobactercloacae complex*** | 25 | 0 | 0 | 1 | 0 | 2 | 0 | 0 | 0 | 0 | 0 | 0 |
| | ***Enterococcus faecalis*** | 0 | 9 | 0 | 0 | 0 | 0 | 0 | 0 | 0 | 0 | 0 | 0 |
| | ***Enterococcus faecium*** | 0 | 0 | 10 | 0 | 0 | 0 | 0 | 0 | 0 | 0 | 0 | 0 |
| | ***Escherichia coli*** | 0 | 0 | 0 | 25 | 0 | 5 | 0 | 1 | 0 | 0 | 0 | 0 |
| | ***Haemophilus influenza*** | 0 | 0 | 0 | 0 | 29 | 0 | 0 | 0 | 0 | 0 | 0 | 0 |
| | ***Klebsiella-pneumoniae*** | 2 | 0 | 0 | 1 | 0 | 28 | 1 | 1 | 0 | 0 | 1 | 1 |
| | ***Pseudomonas aeruginosa*** | 0 | 0 | 1 | 0 | 0 | 0 | 53 | 1 | 2 | 0 | 0 | 0 |
| | ***Staphylococcus aureus*** | 0 | 0 | 0 | 0 | 0 | 0 | 0 | 220 | 5 | 0 | 1 | 0 |
| | ***Staphylococcus epidermidis*** | 0 | 0 | 0 | 0 | 0 | 0 | 1 | 7 | 25 | 0 | 0 | 0 |
| | ***Streptococcus agalactiae*** | 0 | 0 | 0 | 0 | 0 | 0 | 0 | 0 | 0 | 19 | 0 | 0 |
| | ***Streptococcus pneumoniae*** | 0 | 0 | 0 | 0 | 0 | 0 | 0 | 1 | 2 | 0 | 17 | 0 |
| | ***Streptococcus pyogenes*** | 0 | 0 | 0 | 0 | 0 | 0 | 0 | 0 | 0 | 0 | 0 | 28 |

**Figure S12.** Log-ratio lasso-based discrimination of species classification on the training set. Recall rates: *E. cloacae complex* 89.29%, *E. faecalis* 100.00%, *E. faecium* 100.00%, *E. coli* 80.65%, *H. influenzae* 100.00%, *K. pneumoniae* 80.00%, *P. aeruginosa* 92.98%, *S. aureus* 97.35%, *S. epidermidis* 75.76%, *S. agalactiae* 100.00%, *S. pneumoniae* 85.00%, *S. pyogenes* 100.00%. Overall accuracy: 92.95%.

**Test Set Confusion Matrix**

| | | Predict | | | | | | | | | | | |
|---|---|---|---|---|---|---|---|---|---|---|---|---|---|
| | Species | *Enterobactercloacae complex* | *Enterococcus faecalis* | *Enterococcus faecium* | *Escherichia coli* | *Haemophilus influenza* | *Klebsiellapneumoniae* | *Pseudomonas aeruginosa* | *Staphylococcus aureus* | *Staphylococcus epidermidis* | *Streptococcus agalactiae* | *Streptococcus pneumoniae* | *Streptococcus pyogenes* |
| True | *Enterobactercloacae complex* | 12 | 0 | 0 | 0 | 0 | 0 | 0 | 0 | 0 | 0 | 0 | 0 |
| | *Enterococcus faecalis* | 0 | 14 | 0 | 0 | 0 | 0 | 0 | 0 | 0 | 0 | 0 | 0 |
| | *Enterococcus faecium* | 0 | 0 | 12 | 0 | 0 | 0 | 0 | 0 | 0 | 0 | 0 | 0 |
| | *Escherichia coli* | 0 | 1 | 0 | 14 | 0 | 0 | 0 | 0 | 0 | 0 | 0 | 0 |
| | *Haemophilus influenza* | 0 | 0 | 0 | 0 | 24 | 0 | 0 | 0 | 0 | 0 | 0 | 0 |
| | *Klebsiella-pneumoniae* | 0 | 0 | 0 | 0 | 0 | 4 | 0 | 0 | 0 | 0 | 0 | 0 |
| | *Pseudomonas aeruginosa* | 0 | 0 | 0 | 0 | 0 | 0 | 4 | 0 | 0 | 0 | 0 | 0 |
| | *Staphylococcus aureus* | 0 | 0 | 0 | 0 | 0 | 0 | 0 | 98 | 0 | 0 | 0 | 0 |
| | *Staphylococcus epidermidis* | 0 | 0 | 0 | 0 | 0 | 0 | 0 | 6 | 8 | 0 | 0 | 0 |
| | *Streptococcus agalactiae* | 0 | 0 | 0 | 0 | 0 | 0 | 0 | 1 | 0 | 7 | 0 | 0 |
| | *Streptococcus pneumoniae* | 0 | 0 | 0 | 0 | 0 | 0 | 0 | 0 | 1 | 0 | 7 | 0 |
| | *Streptococcus pyogenes* | 0 | 0 | 0 | 0 | 0 | 0 | 0 | 0 | 0 | 0 | 0 | 12 |

**Figure S13.** Log-ratio lasso-based discrimination of species classification on the test set. Recall rates: *E. cloacae* complex 75.00%, *E. faecalis* 75.00%, *E. faecium* 100.00%, *E. coli* 92.86%, *H. influenzae* 100.00%, *K. pneumoniae* 80.00%, *P. aeruginosa* 100.00%, *S. aureus* 100.00%, *S. epidermidis* 66.67%, *S. agalactiae* 100.00%, *S. pneumoniae* 100.00%, *S. pyogenes* 100.00%. Overall accuracy: 94.35%.

**Test Forest Validation Set Confusion Matrix**

| | | Predict | | | | | | | | | | | |
|---|---|---|---|---|---|---|---|---|---|---|---|---|---|
| | **Species** | *Enterobactercloacae complex* | *Escherichia coli* | *Haemophilus influenza* | *Klebsiellapneumoniae* | *Pseudomonas aeruginosa* | *Enterococcus faecalis* | *Enterococcus faecium* | *Staphylococcus aureus* | *Staphylococcus epidermidis* | *Streptococcus agalactiae* | *Streptococcus pneumoniae* | *Streptococcus pyogenes* |
| True | *Enterobactercloacae complex* | 8 | 2 | 0 | 1 | 1 | 0 | 0 | 0 | 0 | 0 | 0 | 0 |
| | *Escherichia coli* | 1 | 12 | 0 | 1 | 0 | 0 | 0 | 0 | 0 | 0 | 0 | 0 |
| | *Haemophilus influenza* | 0 | 0 | 11 | 0 | 1 | 0 | 0 | 0 | 0 | 0 | 0 | 0 |
| | *Klebsiella-pneumoniae* | 0 | 2 | 0 | 13 | 0 | 0 | 0 | 0 | 0 | 0 | 0 | 0 |
| | *Pseudomonas aeruginosa* | 0 | 0 | 0 | 0 | 24 | 0 | 0 | 0 | 0 | 0 | 0 | 0 |
| | *Enterococcus faecalis* | 0 | 0 | 0 | 0 | 0 | 4 | 0 | 0 | 0 | 0 | 0 | 0 |
| | *Enterococcus faecium* | 0 | 0 | 0 | 0 | 0 | 0 | 4 | 0 | 0 | 0 | 0 | 0 |
| | *Staphylococcus aureus* | 0 | 0 | 0 | 0 | 0 | 0 | 0 | 102 | 0 | 0 | 0 | 0 |
| | *Staphylococcus epidermidis* | 0 | 0 | 0 | 0 | 0 | 0 | 0 | 3 | 12 | 0 | 0 | 0 |
| | *Streptococcus agalactiae* | 0 | 0 | 0 | 0 | 0 | 0 | 0 | 0 | 0 | 8 | 0 | 0 |
| | *Streptococcus pneumoniae* | 0 | 0 | 0 | 1 | 0 | 0 | 0 | 0 | 0 | 0 | 7 | 0 |
| | *Streptococcus pyogenes* | 0 | 0 | 0 | 0 | 0 | 0 | 0 | 0 | 0 | 0 | 0 | 12 |

**Figure S14.** Random forest classification of species on the test set. Recall rates: *E. cloacae* complex 66.67%, *E. coli* 85.71%, *H. influenzae* 91.67%, *K. pneumoniae* 86.67%, *P. aeruginosa* 100.00%, *E. faecalis* 100.00%, *E. faecium* 100.00%, *S. aureus* 100.00%, *S. epidermidis* 80.00%, *S. agalactiae* 100.00%, *S. pneumoniae* 87.50%, *S. pyogenes* 100.00%. Overall accuracy: 94.35%.

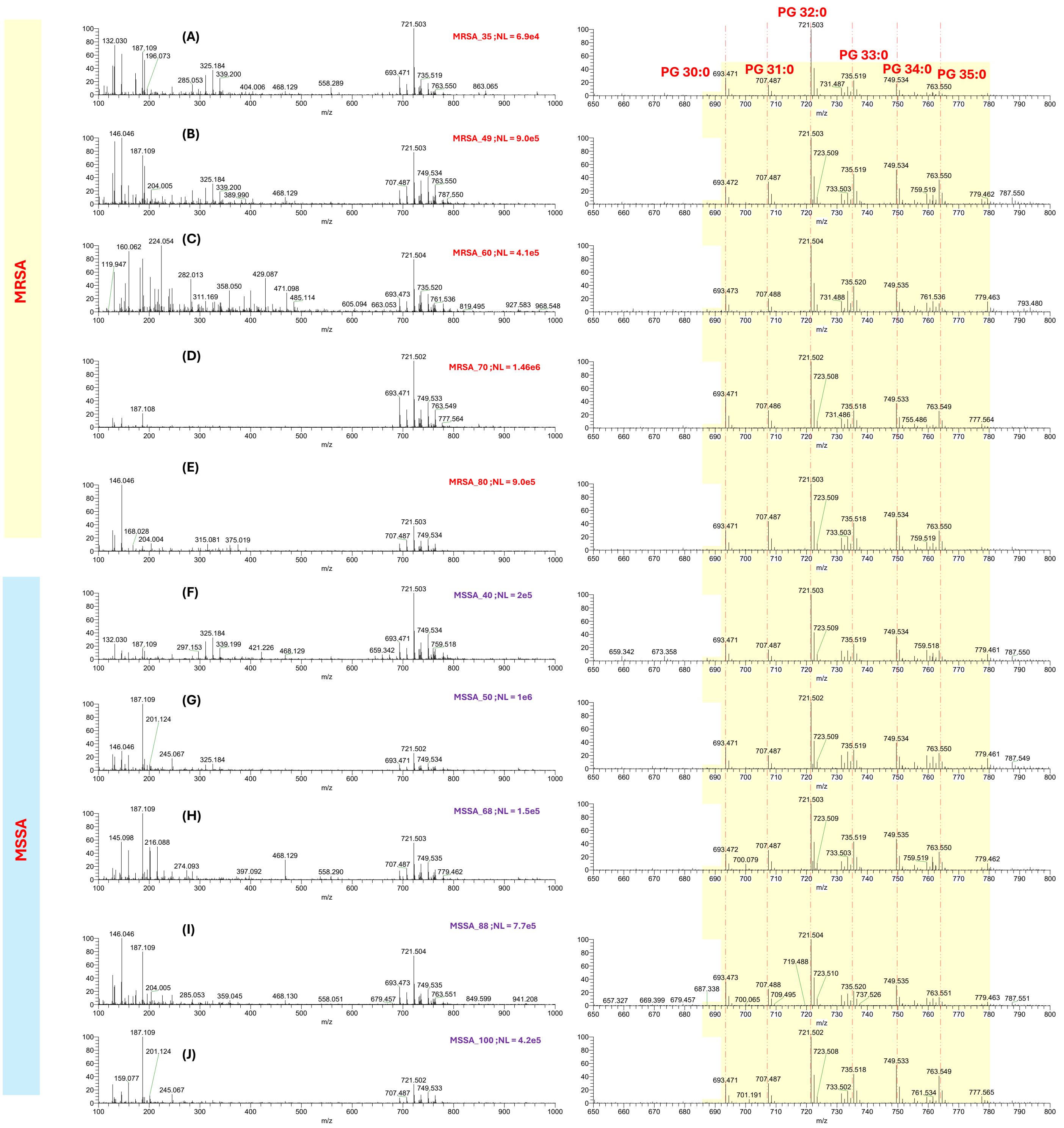

**Figure S15.** MSPen spectra of MSSA and MRSA isolates. The left-hand side shows the mass range of *m/z* 100–1000, while the right-hand side shows the zoomed-in MSPen spectra in the mass range of *m/z* 650–800 (signature range for bacterial species). Panels A–E represent five randomly selected MRSA isolates, and panels F–J represent five randomly selected MSSA isolates, all grown on blood agar. The signature m/z range for *S. aureus* (650–800 Da) is shown on the left-hand side of the spectra for all 10 randomly selected isolates (5 MRSA and 5 MSSA). The MSPen spectra highlights the major membrane lipids, which predominantly belong to the phosphatidylglycerol (PG) class and are annotated above the corresponding *m/z* values. In terms of relative intensity, with respect to *m/z* 721.504 (PG 32:0), all spectra appear highly similar within the *m/z* 650–800 range. This high similarity in lipid profiles may be one of the reasons for the poor performance of the statistical classifiers. In future work, we plan to increase the number of MSSA and MRSA isolates to improve discrimination between the two groups.

A)

| Training Set | | Predict | |
|---|---|---|---|
| | Antibiotic Resistance | MRSA | MSSA |
| True | MRSA | 66 | 36 |
| | MSSA | 34 | 90 |

| Class | Recall Rate |
|---|---|
| MRSA: | 64.7% |
| MSSA: | 72.6% |
| Overall | 69.0% |

B)

| Test Set | | Predict | |
|---|---|---|---|
| | Antibiotic Resistance | MRSA | MSSA |
| True | MRSA | 31 | 25 |
| | MSSA | 9 | 37 |

| Class | Recall Rate |
|---|---|
| MRSA: | 55.4% |
| MSSA: | 80.4% |
| Overall | 66.7% |

**Figure S16.** Logistic regression lasso discrimination of MRSA versus MSSA on the **(A)** training set and **(B)** test set.

A)

| Training Set | | Predict | |
|---|---|---|---|
| | Antibiotic Resistance | MRSA | MSSA |
| True | MRSA | 61 | 41 |
| | MSSA | 44 | 80 |

| Class | Recall Rate |
|---|---|
| MRSA: | 59.8% |
| MSSA: | 64.5% |
| Overall | 62.4% |

B)

| Test Set | | Predict | |
|---|---|---|---|
| | Antibiotic Resistance | MRSA | MSSA |
| True | MRSA | 32 | 24 |
| | MSSA | 15 | 31 |

| Class | Recall Rate |
|---|---|
| MRSA: | 57.1% |
| MSSA: | 67.4% |
| Overall | 61.8% |

**Figure S17.** Log-ratio lasso discrimination of MRSA versus MSSA on the **(A)** training set and **(B)** test set.

| Test Set | | Predict | |
|---|---|---|---|
| | Antibiotic Resistance | MRSA | MSSA |
| True | MRSA | 26 | 30 |
| | MSSA | 5 | 41 |

| Class | Recall Rate |
|---|---|
| MRSA: | 46.4% |
| MSSA: | 89.1% |
| Overall | 65.7% |

**Figure S18.** Random forest classification of MRSA versus MSSA on test set.

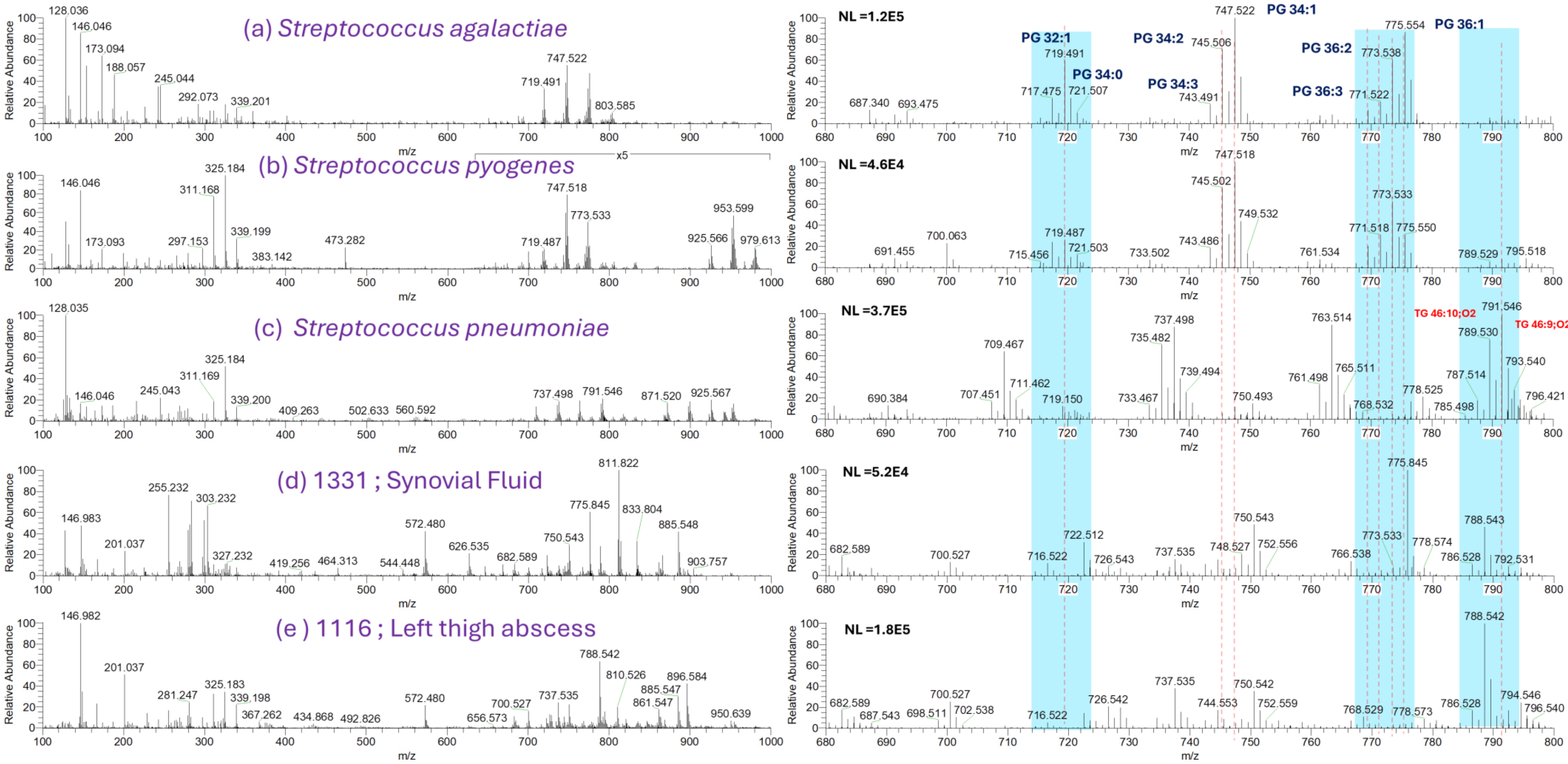

**Figure S19.** MSPen spectra of *Streptococcus* species and infected samples: (a) *S. agalactiae*, (b) *S. pyogenes*, (c) *S. pneumoniae*, (d) Synovial fluid (*S. pyogenes*, 4+; sample 1331) and (e) Left thigh abscess fluid (*S. agalactiae*, 4+; sample 1116). The infected tissue (sample 1331) exhibited characteristic peaks at *m/z* 745.501, *m/z* 747.518, *m/z* 769.501, *m/z* 771.515, and *m/z* 773.533, consistent with a monomicrobial S. pyogenes infection. Sample 1116 displayed peaks at *m/z* 745.508, *m/z* 747.518, *m/z* 771.515, and *m/z* 773.533, indicating the presence of *Streptococcus* species.

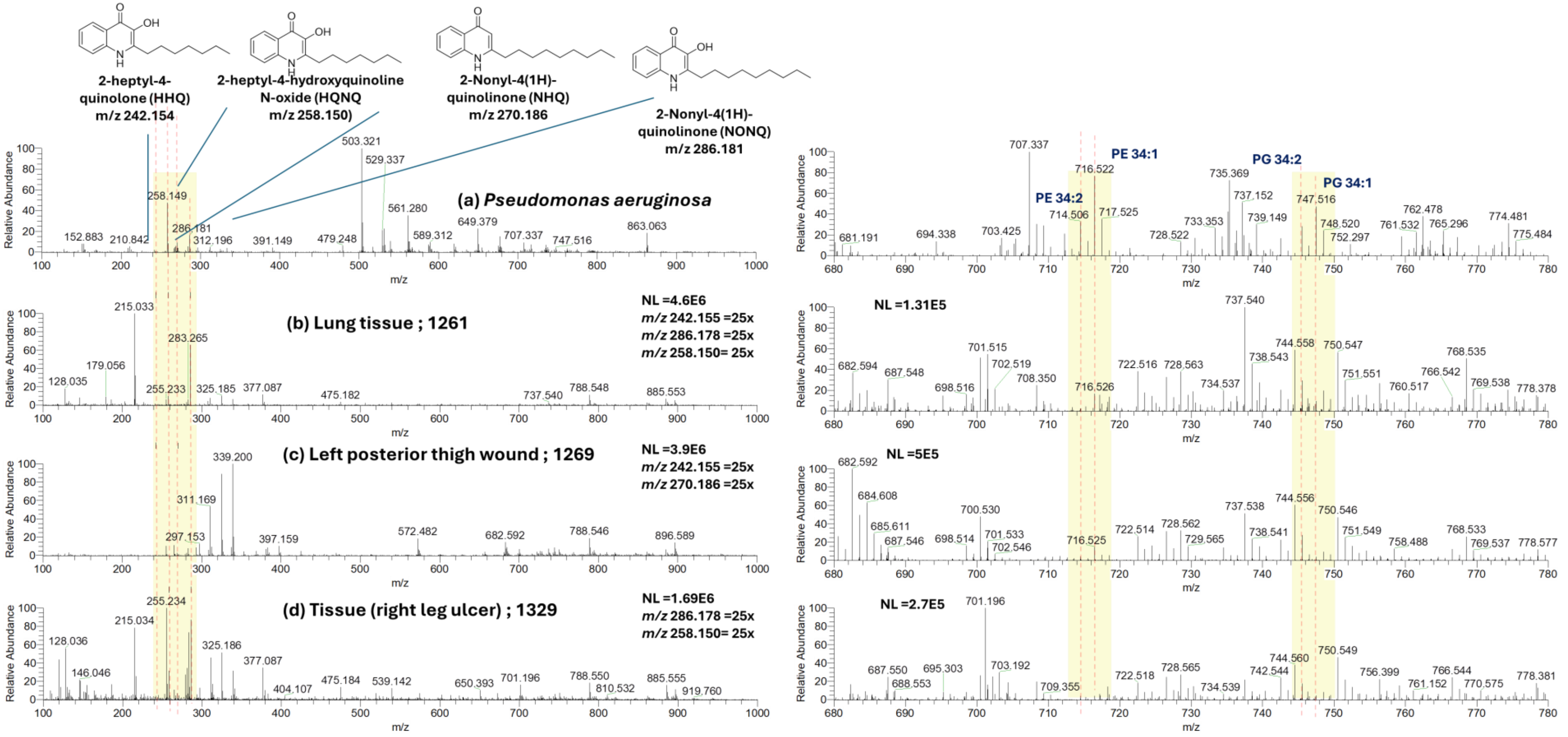

**Figure S20.** MSPen spectra of *P. aeruginosa*-infected samples. (a) Clinical isolate of *P. aeruginosa*; (b) Lung tissue sample (*P. aeruginosa* 4+; sample 1261); (c) Left posterior thigh wound (*P. aeruginosa* and *P. mirabilis* (4+); *K. oxytoca,* and *S. oralis* (2+); and *K. variicola* (1+) ; sample 1269) and (d) tissue (right leg ulcer, *P. aeruginosa* 4+; sample 1329). The left-hand side of each spectrum shows the detection range of *m/z* 100–1000, while the right-hand side presents a zoomed-in view of the *m/z* 680–780 region. The tissue samples were infected with *P. aeruginosa*, exhibiting a 4+ level of infection. In the MSPen spectra of the corresponding pure isolate, the *m/z* 200–400 region corresponds to quorum sensing (QS) molecules and the *m/z* 700–800 range to bacterial membrane lipids. In sample 1261, three PQS-related molecules were detected: HHQ (*m/z* 242.150), HQNO (*m/z* 258.150), and NQNO (*m/z* 286.181). In sample 1269, HHQ (*m/z* 242.155) and db-NHQ (*m/z* 270.186) were detected. In sample 1329, HQNO (*m/z* 258.150) and NQNO (*m/z* 286.181) were observed. Additionally, lipid peaks were observed in the *m/z* range of 710–720, with *m/z* 714.509 and *m/z* 716.522 tentatively assigned to PE 34:2 and PE 34:1, respectively. However, as these lipid species are also commonly present in human tissue, PQS-related molecules were selected as more specific molecular identifiers of *Pseudomonas aeruginosa* infection.

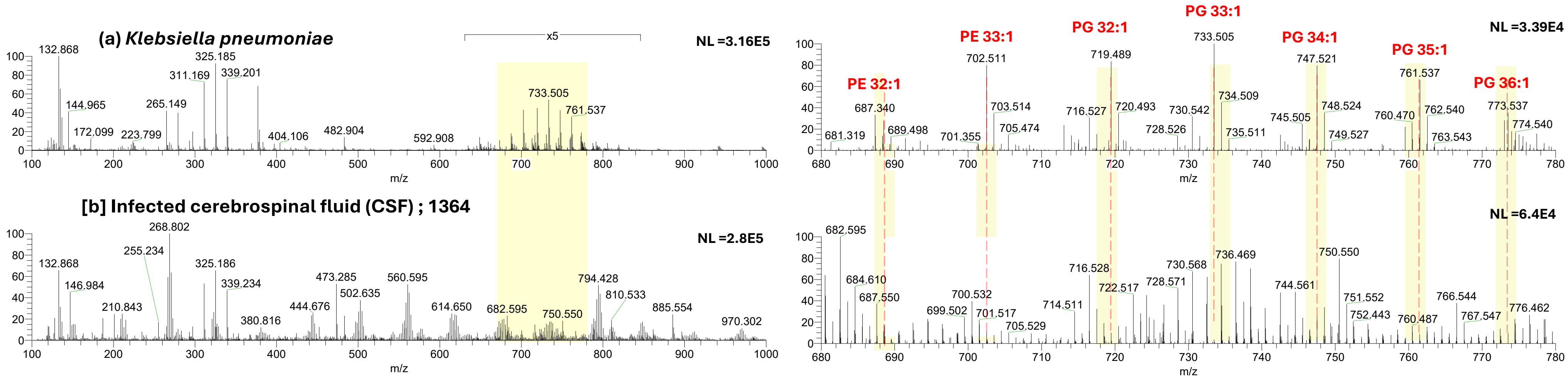

**Figure S21.** MSPen spectra of Infected cerebral spinal fluid (4+). (a) Full mass range spectrum (*m/z* 100–1000); of pure isolate of *K. pneumoniae* (b) Full mass range spectrum (*m/z* 100–1000); of Infected CSF, the right-hand side shows the full MS view (*m/z* 100–1000), while the left-hand side displays an expanded view of the *m/z* 680–780 region. On the right-hand side, the *m/z* 650–780 region representing the signature range for bacterial species is highlighted in light pink. On the left-hand side, bacterial-specific peaks are highlighted in light blue, with a dotted red line indicating the presence of a bacterial-specific peak in the infected sample. In the MSPen analysis of cerebrospinal fluid (CSF), the sample initially appeared transparent and exhibited a strong salt signal, likely due to its ionic content. To reduce this interference, the sample was centrifuged at 20,000 rpm for 5 minutes, yielding a small white pellet. The pellet was washed twice with 200 µL of water, vortexed, and centrifuged to remove residual salts. Finally, 200 µL of methanol was added before proceeding with MSPen analysis, Key bacterial lipids identified include *m/z* 702.5079 (PE 33:1), *m/z* 719.490 (PG 32:2), *m/z* 733.503 (PG 33:1), *m/z* 747.518 (PG 34:1) and *m/z* 773.537 (PG 36:2).

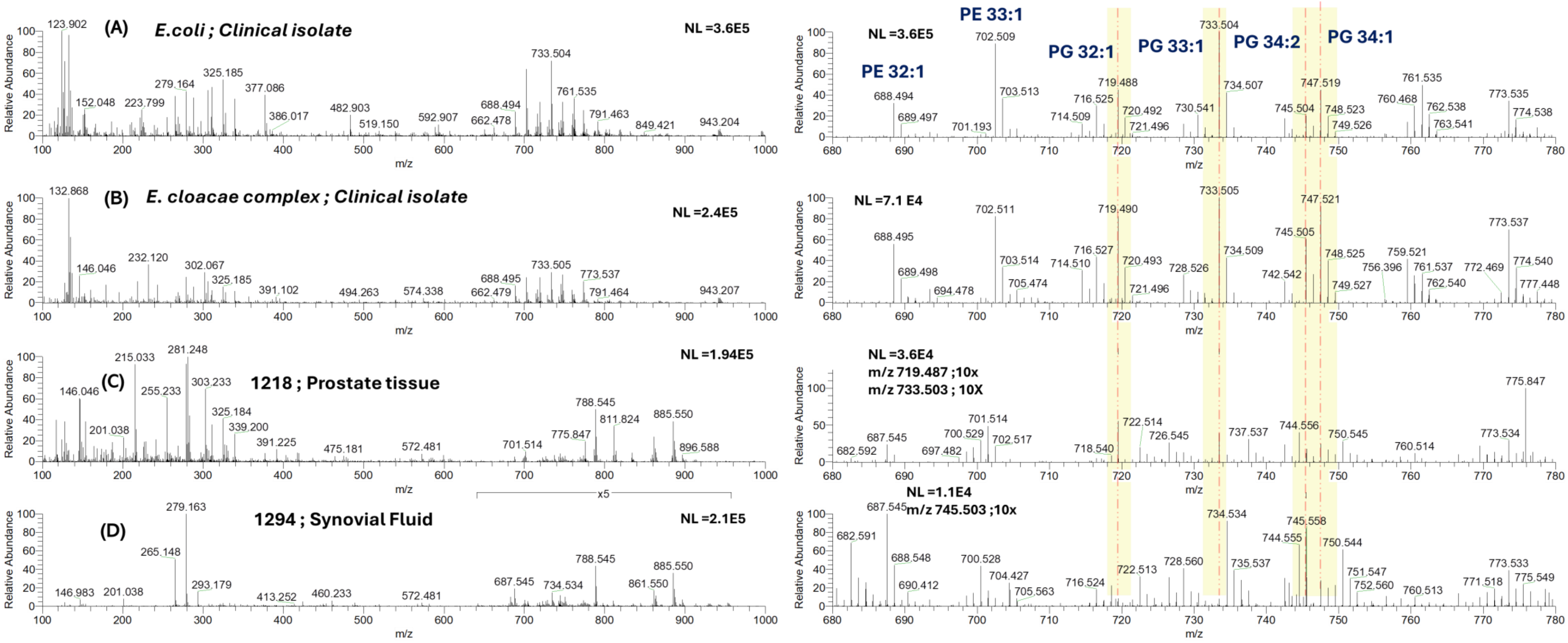

**Figure S22**. MSPen spectra of specimens infected with members of the Enterobacteriaceae family. MSPen spectra of infected specimens showing (A) the clinical isolate of *Escherichia coli*, (B) the clinical isolate of *Enterobacter cloacae* complex, (C) infected prostate tissue (*E. coli*, 4+; sample 1218), and (D) infected synovial fluid (*Enterobacter cloacae* complex, 4+, sample 1294). As shown in panels (A) and (B), prominent peaks at *m/z* 702.508, *m/z* 719.490, *m/z* 733.508, *m/z* 745.504, and *m/z* 747.518 were observed in pure isolates of *E. coli* and the *E. cloacae* complex. In infected prostate tissue (sample1218), peaks at *m/z* 719.490, *m/z* 733.505, *m/z* 745.505, and *m/z* 747.518 were observed. In the synovial fluid sample (sample 1294), peaks at *m/z* 745.505 and *m/z* 747.518 were detected, consistent with the pattern seen in *E. cloacae* complex isolates.

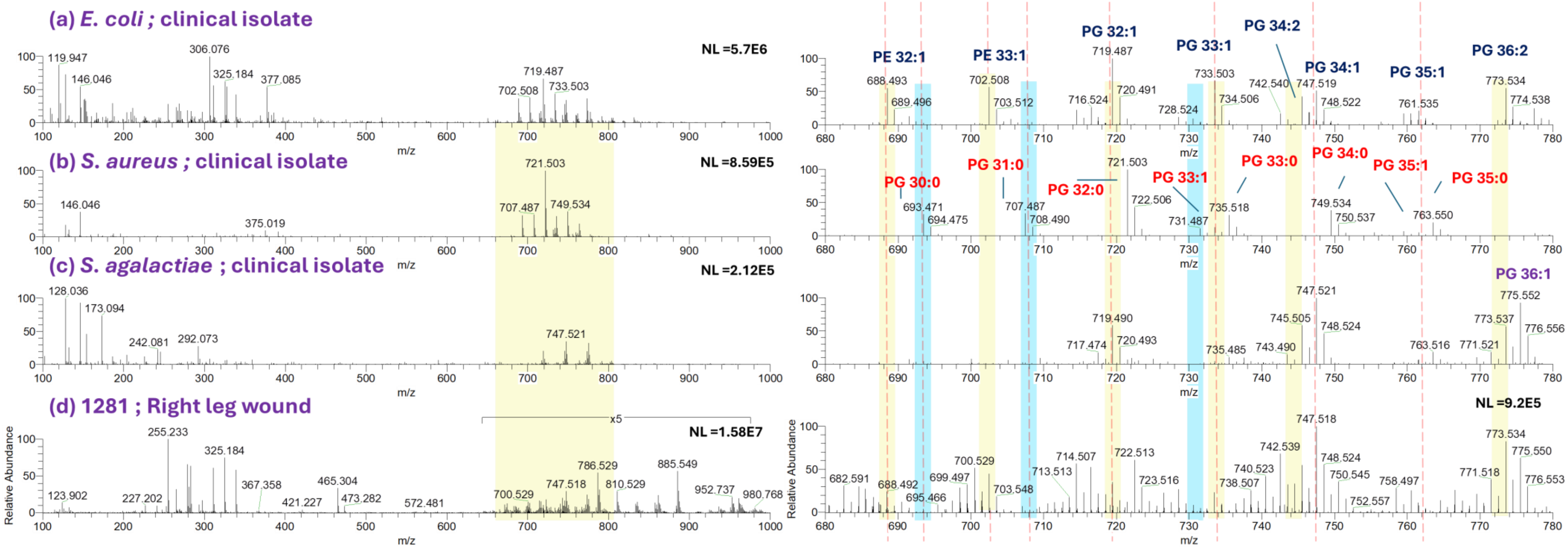

**Figure S23.** MSPen spectra (*m/z* 100 – 1000; zoomed in *m/z* 680 – 780) of polymicrobial-infected tissues and gram-positive and gram-negative clinical isolates. Shown are: (a*) E. coli* (clinical isolate), (b) *S. aureus* (clinical isolate), (c) *S. agalactiae* (clinical isolate), and (d) right leg wound (*K. pneumoniae* 4+, *K. oxytoca* 4+, *M. morganii* 2+, *S. aureus* 3+, *E. faecalis* 3+, *P. mirabilis* 4+, *V. fluvialis* 1+, *P. somerae* 4+; sample 1281).

Sample 1281: (right leg wound) contained *K. pneumoniae*, *K. oxytoca*, *P. mirabilis*, and *P. somerae* (4+); *S. aureus* and *E. faecalis* (3+); *M. morganii* (2+); and *V. fluvialis* (1+). MS-Pen spectra revealed *m/z* 693.471, *m/z* 707.487, *m/z* 721.502 (gram positive) and *m/z* 688.489, *m/z* 702.508, *m/z* 719.488, *m/z* 733.503 (gram negative), confirming polymicrobial infection, though species-level differentiation was limited due to overlapping lipid features.

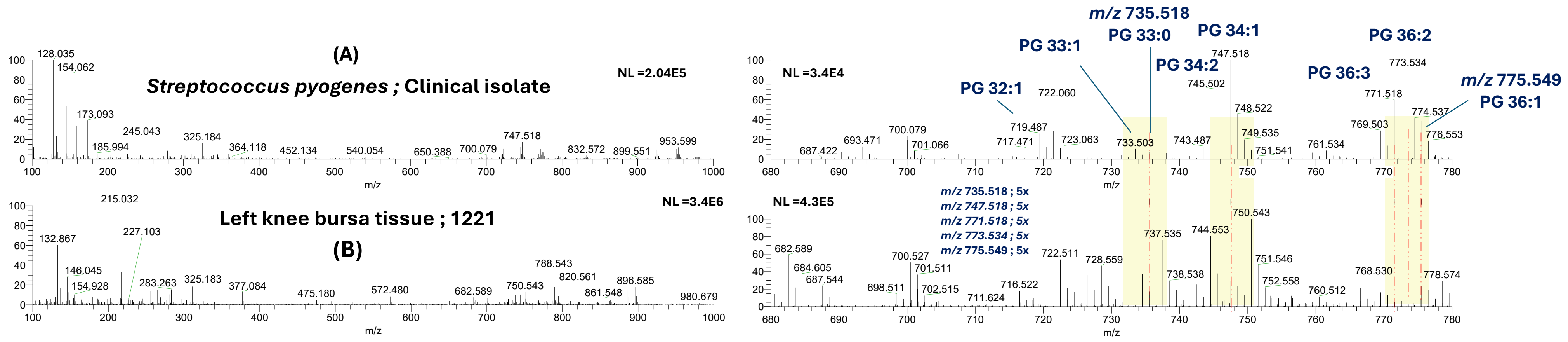

**Figure S24.** MSPen spectra (*m/z* 100 – 1000; zoomed in *m/z* 680 – 780) of polymicrobial-infected tissues and gram-positive clinical isolates. Shown are: (a*) S. pyogenes* (clinical isolate) and (b) left knee bursa tissue (sample 1221*, S. pyogenes (4+) and S. aureus (1+).* In the infected sample, we observed features at m/z 735.518 (PG 33:0), 745.502 (PG 34:2), 747.518 (PG 34:1), 771.518 (PG 36:3), 773.534 (PG 36:2), and 775.549 (PG 36:1), which match the spectral profile of *S. pyogenes* clinical isolate.

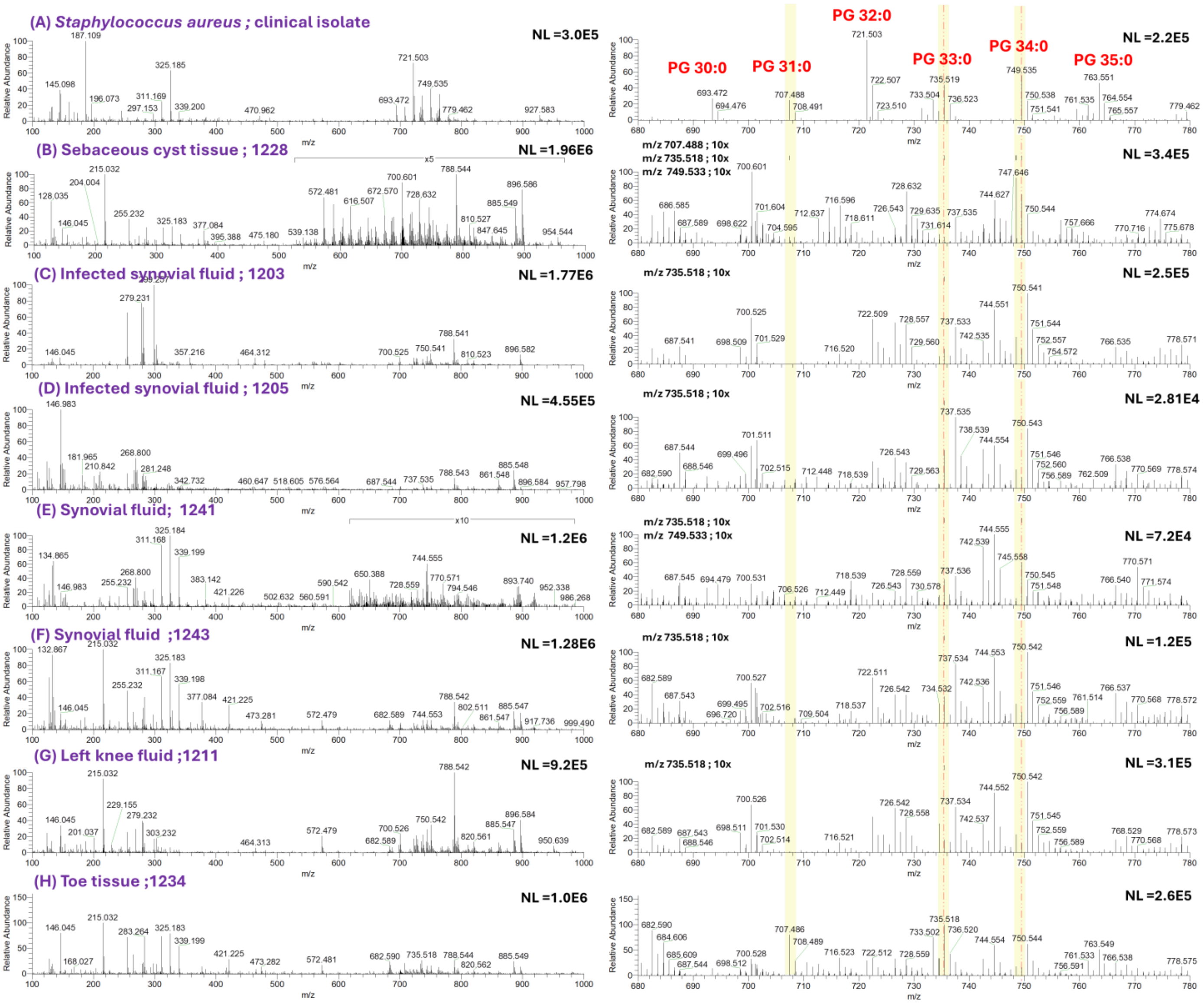

**Figure S25.** MSPen spectra of *S. aureus* and infected clinical (*S. aureus*; 4+) are shown. The left panel displays the m/z range of 100–1000, and the right panel shows a zoomed-in view of the 680–780 m/z range. (A) *S. aureus*, clinical isolate; (B) sebaceous cyst tissue, (sample 1228); (C) infected synovial fluid (sample 1203); (D) infected synovial fluid (sample 1205); (E) synovial fluid (sample 1241); (F) synovial fluid (sample 1243); (G) left knee fluid (sample 1211); and (H) toe tissue (sample 1234). The pure clinical *S. aureus* isolate shows characteristic peaks at *m/z* 693.471 (PG 30:0), *m/z* 707.485 (PG 31:0), *m/z* 721.505 (PG 32:0), *m/z* 735.521 (PG 33:0), *m/z* 749.549 (PG 34:0), and *m/z* 763.549 (PG 35:0), which were annotated in the pure clinical isolate. In the overlay of the infected clinical specimens, the peaks at *m/z* 735.521 (PG 33:0) and *m/z* 749.549 (PG 34:0) were commonly observed and are indicated with a fine dotted red line, corresponding to *S. aureus*. Interestingly, in sample code 1234 (infected toe), all the major m/z peaks of *S. aureus* were detected.

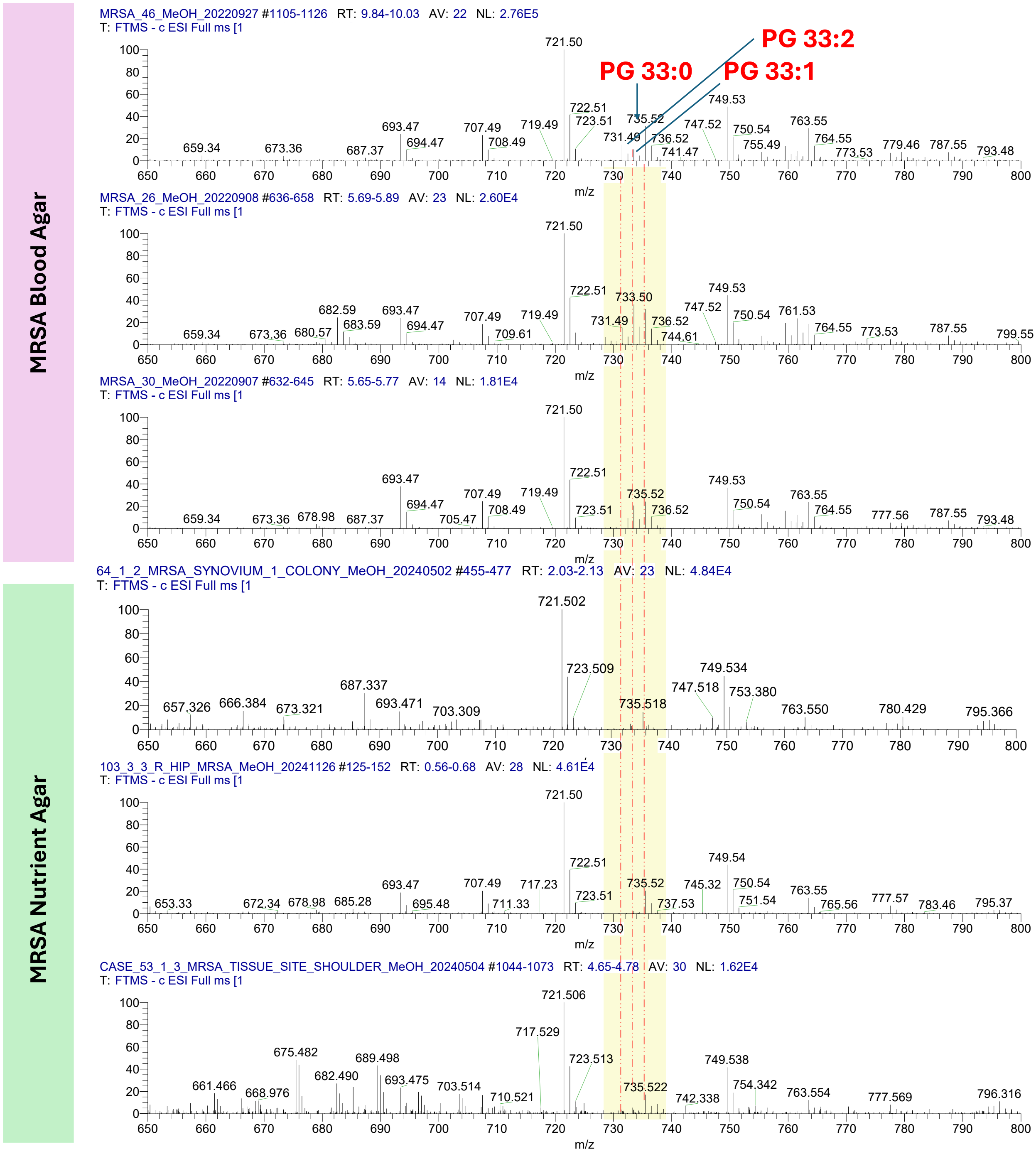

**Figure S26**. Zoomed-in mass spectra of MRSA grown on nutrient agar and blood agar. The top three spectra correspond to *S. aureus* grown on blood agar, while the bottom three correspond to cultures grown on nutrient agar. Blood agar contains 5% sheep red blood cells, making it more nutrient-rich than nutrient agar, which lacks red blood cells. A clear difference between the two media is observed in the *m/z* 731–737 range, where PG 33:2 (*m/z* 731.497), PG 33:1 (*m/z* 733.504), and PG 33:0 (*m/z* 735.518) are clearly detected in blood agar. In contrast, the peaks corresponding to PG 33:2 (*m/z* 731.497) and PG 33:1 (*m/z* 733.504) show relatively lower intensity in nutrient agar.

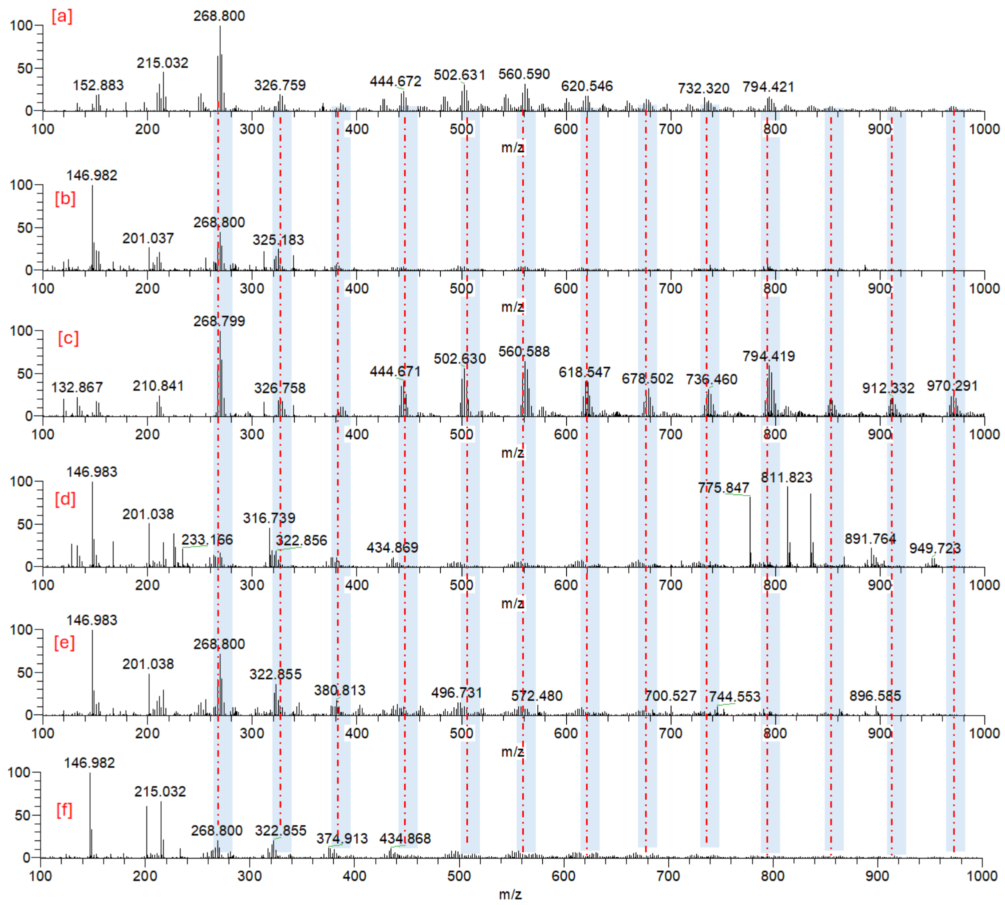

**Figure S27.** MS-Pen spectra in negative ion mode (*m/z* 100–1000) from various infected specimens, showing characteristic salt (NaCl) signatures: (a) right knee prosthetic joint infection (PJI; *S. mitis* 4+, sample 1199), (b) left knee tissue (*S. aureus* 4+, sample 1204), (c) right foot tissue from a diabetic ulcer (S. epidermidis 4+, sample 1268), (d) right hip tissue (*S. aureus* [MSSA] 4+, sample 1289), (e) cerebrospinal fluid (CSF; *E. coli* 4+, sample 1293), and (f) right knee PJI (*S. aureus* 4+, sample 1208). The peak at *m/z* 268.800 $[M+Cl]^-$, where M = $(NaCl)_4$, was followed by a series of clusters showing an incremental increase of 57.985 Da per additional NaCl molecule, extending up to *m/z* 970.291. Bacterial and human tissue lipids were predominantly observed in the range of *m/z* 650–900, where the presence of NaCl cluster ions limited further analysis.

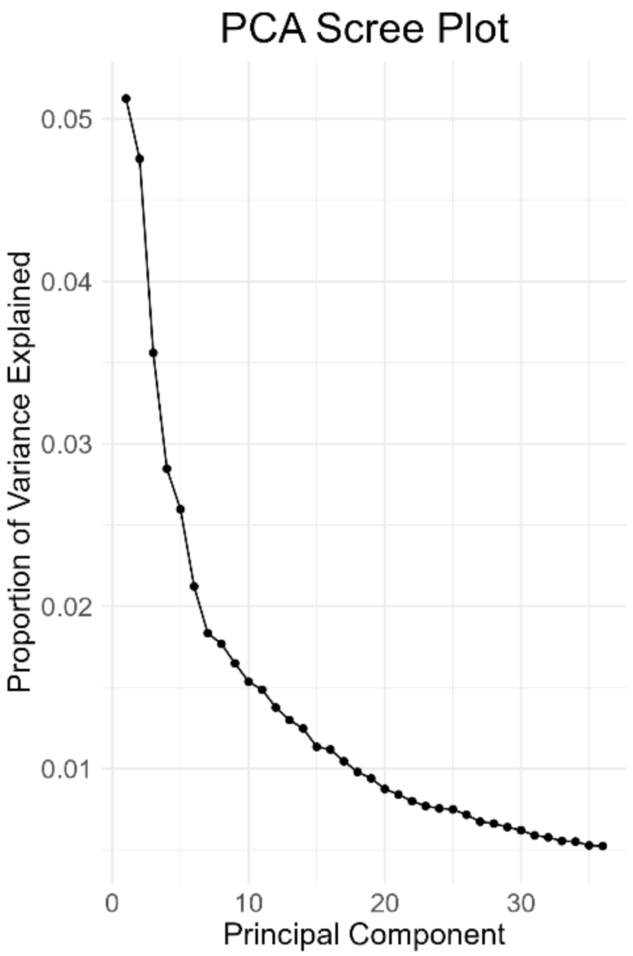

**Figure S28.** Scree plot of the principal components from PCA analysis on the x-axis, with the respective variance explained over the data on the y-axis.

**Table S1.** Bacterial growth and MALDI results.

| | Sample code | Name of Microbes | Infection site | | Bacterial Growth | MALDI results | Confidence level |
|---|---|---|---|---|---|---|---|
| 1 | 1063 | Coagulase-negative staphylococci | Right hip PJI | Adult | 2+ | S. Epidermidis | 99.9 |
| 2 | 1275 | *S. aureus* (MSSA) | Right elbow bursa | | 4+ | *S. aureus* | 99.9 |
| 3 | 1277 | *S. aureus* (MSSA) | Left hip | Adult | 2+ | *S. aureus* | 99.9 |
| 4 | 1359 | *S. aureus* (MRSA) | Thora fluid | | 1+ | *S. aureus* | 99.9 |
| 5 | 1046 | *S. aureus* (MSSA) | Breast Abscess | | 4+ | *S. aureus* | 99.9 |

### Supplemental Methods:

Desalting and Centrifugation

Samples such as pus, tiny tissue fragments suspended in saline, and cerebrospinal fluid are rich in salt which causes ion suppression when analyzed with a mass spectrometer. To remove salt, 200 μL of the sample suspension was centrifuged 20,000 rpm for five minutes at 4°C. The supernatant was discarded, and 200 μL of LC-MS grade water was added. After vertexing for 30 seconds, the sample was centrifuged again under the same conditions. This washing step was repeated twice. The final pellet was resuspended in 200 μL of LC-MS grade methanol and centrifuged at 10,000 rpm. Finally, 50 μL of the supernatant was aspirated and analyzed using the MS Pen source.
